# tRNA thiolation defects disrupt cellular proteostasis and tissue homeostasis in mammals

**DOI:** 10.1101/2025.10.24.684405

**Authors:** Lukas Englmaier, Marta Walczak, Daniel Malzl, Cristian Eggers, Felix Eichin, Valentina C. Sladky, Filip M. Gallob, Yevheniia Tyshchenko, Zarghun Zarif, Thomas Kolbe, Lama Al-Abdi, Jörg Menche, Nathalie Spassky, Stephan Geley, Walter Rossmanith, Fowzan S. Alkuraya, Sebastian A. Leidel, Sebastian Glatt, Andreas Villunger

## Abstract

Sulfur modification of tRNA wobble uridines is an evolutionarily conserved mechanism that ensures efficient protein synthesis. In humans, loss of this anticodon modification due to mutations in *CTU2* (cytosolic thiouridylase 2) causes DREAM-PL syndrome, a severe congenital disorder often leading to early postnatal death. However, the mechanisms by which loss of tRNA thiolation drives pathology remain unclear. Here, we show that loss of *CTU2* triggers significant cellular proteostasis defects in patient cells and model cell lines. Structural and biochemical analyses reveal that the pathogenic CTU2^L63P^ mutation destabilizes the CTU1/CTU2 complex and abolishes tRNA binding and thiolation. Acute loss of *CTU2* caused codon-specific ribosome pausing at A-ending codons decoded by thiolated tRNAs, and decreased ribosome occupancy of A-rich transcripts in a dosage-dependent manner. Codon-biased mRNAs transcribed from genes critical for ciliogenesis are predicted to be most affected, linking their reduced translation to DREAM-PL etiology in humans. Surprisingly, *Ctu2^L63P^* mice display severe thiolation defects, but develop normally, are viable and fertile. Our findings highlight the importance of functional tRNA thiolation for organismal health in humans and identify species-specific vulnerabilities during embryonic development in mammals.

## INTRODUCTION

Transfer RNAs (tRNAs) ensure accurate translation of messenger RNA (mRNA) into protein. Although approximately 430 human tRNA genes are predicted in the human genome (Chan & Lowe, 2016), only a subset is actually expressed. Because the expressed tRNA anticodon repertoire does not cover all 61 sense codons used in mRNA (Gao et al., 2024), decoding relies on flexible base pairing at the third, or “wobble,” position of the codon (Crick, 1966). tRNA modifications at the corresponding anticodon position 34, particularly of uridine, expand pairing capacity while maintaining translational accuracy (Agris et al., 2018).

Among these “wobble modifications”, tRNA thiolation is a key regulator of translation conserved across all domains of life. In vertebrates, a complex enzymatic cascade replaces oxygen at the 2-carbon of uracil (U_34_) with sulfur (s^2^) in 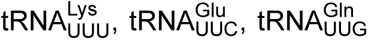 and 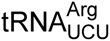 (Björk et al., 2007; Yoshida et al., 2015). This modification is catalyzed by the cytosolic thiouridylase CTU1:CTU2 complex (Dewez et al., 2008) as the final step of the URM1 sulfur relay pathway (Huang et al., 2008; Leidel et al., 2009; Noma et al., 2009; Schlieker et al., 2008). In eukaryotes, thiolation is strongly enhanced when U_34_ is first modified with a methoxycarbonylmethyl group (mcm^5^) that is initiated by the Elongator complex (ELP1-6) and completed by the methyltransferase ALKBH8 (Abbassi et al., 2024; Huang et al., 2005; Songe-Møller et al., 2010). The s² group stabilizes the anticodon loop for cognate U:A pairing, while mcm⁵ facilitates U:G wobbling, allowing mcm⁵s²-modified tRNAs to decode both A- and G-ending codons efficiently (Durant et al., 2005; Vendeix et al., 2012). Together with other anticodon modifications, tRNA thiolation helps maintain the reading frame and prevents mistranslation within mixed codon boxes (Tükenmez et al., 2015).

At the molecular level, 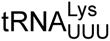 lacking s^2^ at U_34_ binds to AAA codons with lower affinity and is frequently rejected by the ribosome (Ranjan & Rodnina, 2017). Accordingly, the speed of translation is decreased, which is in line with observed ribosome pausing at cognate AAA and CAA codons upon the loss of either mcm^5^ or s^2^ in yeast (Nedialkova & Leidel, 2015; Zinshteyn & Gilbert, 2013). This codon-specific slowdown triggers stress signaling and perturbs proteostasis, which in its most severe form can cause protein loss through aggregation (Nedialkova & Leidel, 2015; Rapino et al., 2021). In yeast, these phenotypes can be alleviated by overexpressing 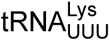 and 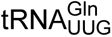 that are normally subject to modification, thereby restoring translation (Björk et al., 2007; Leidel et al., 2009; Nedialkova & Leidel, 2015).

Dependence on tRNA wobble modifications varies across biological systems. Yeast and plants are generally viable without fully modified tRNAs, but mutants show growth defects under stress (Björk et al., 2007; Esberg et al., 2006; Leiber et al., 2010; Leidel et al., 2009; Xu et al., 2020). Likewise, HEK293T cells can tolerate loss of *ELP3* or *CTU2* (Linder et al., 2025). However, the combined loss of mcm^5^ and s^2^ modifications has additive detrimental effects, exemplified by developmental defects in double-mutant worms (C. Chen et al., 2009). In contrast, deletion of the elongator subunit *Elp3* is lethal in fruit flies and mice (Bento-Abreu et al., 2018; Walker et al., 2011). Mutations in human Elongator subunits and other tRNA wobble modifiers lead to severe neurological and developmental disorders, including Familial Dysautonomia (*ELP1*) (Slaugenhaupt et al., 2001), amyotrophic lateral sclerosis (*ELP3*-associated) (Simpson et al., 2009), and neurodevelopmental disorders (*ELP1*, *ELP2*, *ELP3*, *ELP4*, *ELP6, ADAT3 and ALKBH8*) (Alazami et al., 2013; Gaik et al., 2022; Kojic et al., 2021, 2023; Monies et al., 2019; Strug et al., 2009).

Similarly, defects in the tRNA thiolation machinery have been linked to a complex pathology that is often severe and may lead to early postnatal death of affected patients (Shaheen, Al-Salam, et al., 2016; Shaheen, Patel, et al., 2016). Consistent with the essential role of this pathway, multiple URM1 pathway components across many human cell lines are predicted to be indispensable (NFS1, MOCS3, URM1, CTU2) (DepMap, 2024; Tsherniak et al., 2017) and their deletion in mice causes embryonic lethality (*Urm1, Ctu1, Ctu2* – International Mouse Phenotyping Consortium IMPC (Groza et al., 2023)). In affected patients, biallelic *CTU2* loss-of-function mutations manifest in impaired tRNA thiolation at the cellular level (Shaheen, Al-Salam, et al., 2016; Shaheen, Patel, et al., 2016). Most mutations disrupt splice sites of the *CTU2* transcript, leading to frameshifts, premature stop codons and protein degradation (Mahoney et al., 2023; Shaaban et al., 2024; Shaheen et al., 2019). Other pathogenic alterations include C-terminal protein truncations and a leucine-to-proline point mutation (*CTU2^L63P^*) (Helsmoortel et al., 2015; Shaheen et al., 2019). The resulting syndrome was termed DREAM-PL syndrome, as severe cases exhibit congenital anomalies including <u>d</u>ysmorphic facies, <u>re</u>nal agenesis, <u>a</u>mbiguous genitalia, <u>m</u>icrocephaly, <u>p</u>olydactyly and lissencephaly (Shaheen, Al-Salam, et al., 2016). The clinical presentation shares features of ciliopathies, with multi-system involvement of the kidneys, brain, and reproductive organs, and typically associated features like polydactyly and lissencephaly (Waters & Beales, 2011). The underlying cellular and molecular defects explaining the complex pathology remain unknown. Here, we present complementary *in vitro* systems and a patient mutation knock-in mouse model to study DREAM-PL syndrome.

## RESULTS

### CTU2 deficiency in DREAM-PL syndrome disrupts tRNA thiolation, cell growth and proteostasis

To investigate the disease mechanisms, we characterized primary dermal fibroblasts and EBV-transformed B-lymphoblastoid cell lines (LCLs) from DREAM-PL patients carrying mostly bi-allelic splice-site mutations in *CTU2.* We also included LCLs carrying either the *CTU2^L63P^* point mutation or a 4-nucleotide deletion resulting in a C-terminal frameshift **(Table 1)**. All DREAM-PL patient cells showed a drastic reduction in CTU2 protein levels **(Fig. 1a)**. CTU1 and CTU2 form a conserved complex, with CTU1 serving as the catalytic tRNA-modifying subunit (Dewez et al., 2008; Liu et al., 2016). Hence, we tested to which degree the absence of CTU2 functionally impacted tRNA thiolation. To this end, we used ([*N*-acryloylamino]phenyl)mercuric chloride (APM)-containing polyacrylamide gels followed by northern blotting (Igloi, 1988). APM slows the migration of modified tRNAs, causing a detectable shift when probed with radiolabeled tRNA-specific probes. Our analysis confirmed a severe reduction of 2-thiolation of 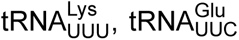 and 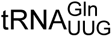 in patient cells, and extends previous findings (Shaheen et al., 2019) by revealing markedly reduced modification of 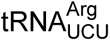 **(Fig. 1b, Suppl. Fig. S1a)**. Importantly, we were able to fully restore tRNA thiolation upon transduction with wildtype *CTU2* **(Fig. 1b)**, confirming that CTU2 is essential for this process in human cells. Interestingly, tRNA thiolation defects varied between fibroblasts and LCLs, as well as among different patient LCLs **(Suppl. Fig. S1b)**.

**Figure 1.**
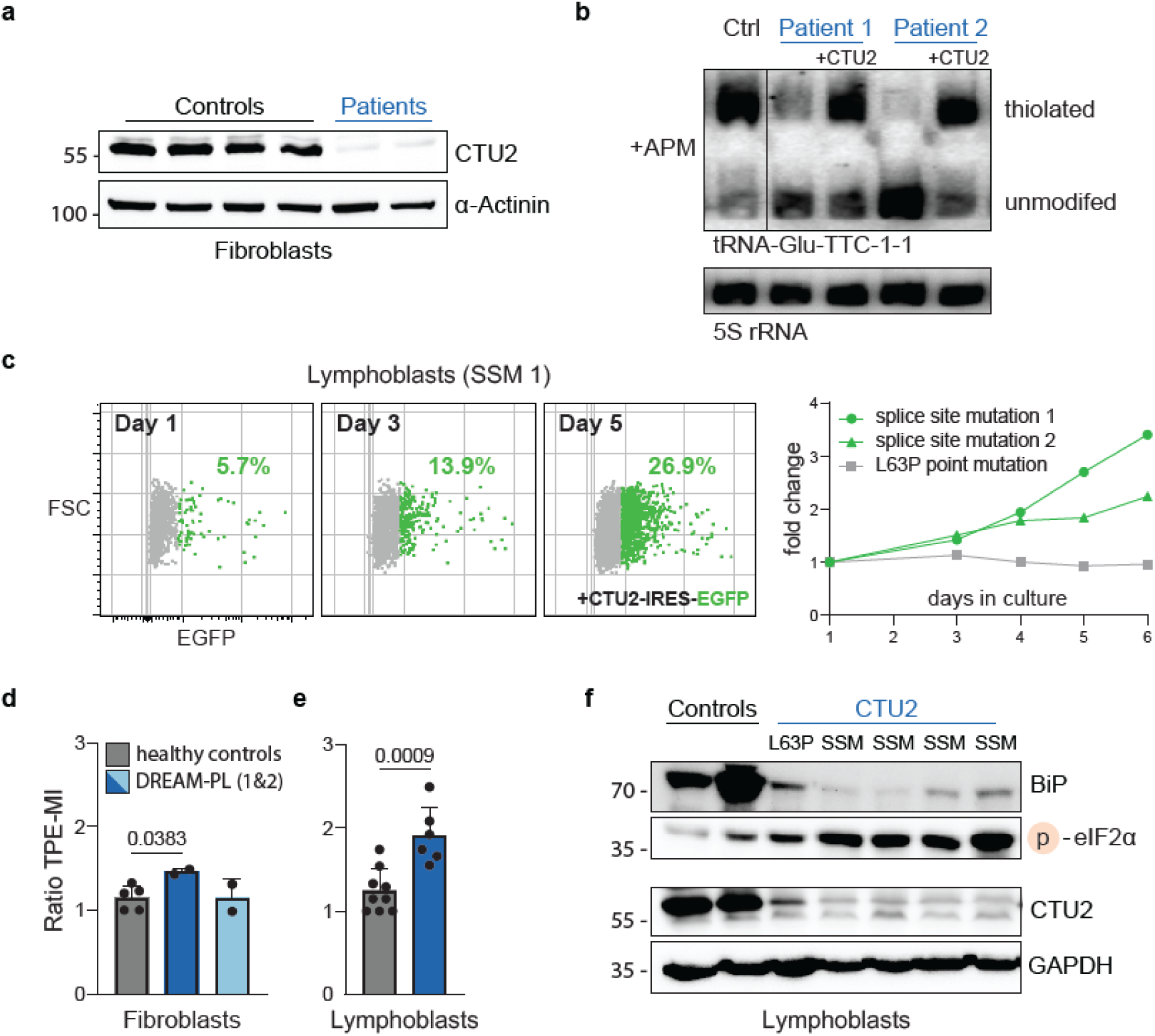
CTU2 deficiency in DREAM-PL disrupts tRNA thiolation, cell growth and proteostasis. **(a)** Western blot analysis of fibroblasts from DREAM-PL patients and controls. **(b)** APM-northern blot in patient fibroblasts. Thiolation is restored upon CTU2 overexpression. 5S rRNA serves as a loading control. **(c)** Growth rescue of patient lymphoblastoid cell lines (LCLs) by CTU2 complementation. Left: CTU2-eGFP–expressing cells (green) outcompete non-transduced cells (grey). Right: comparison of three patient LCLs with the indicated CTU2 mutations (technical replicates). **(d, e)** Quantification of protein unfolding by TPE-MI staining in patient fibroblasts (technical duplicates) and (e) lymphoblasts (n = 3, technical triplicates) analyzed by flow cytometry. Controls from healthy donors are shown in grey. Normalized mean fluorescence intensity reflects TPE-MI binding to exposed cysteines. Data represent mean ± SD; two-tailed unpaired t-tests, p-values are indicated. **(f)** Western blot analysis of proteostasis-related markers in patient LCLs. SSM, splice-site mutation.

**Table 1.** DREAM-PL patient derived cells.

| ID | DNA sequence | Protein sequence | Cell type | Description | Reference |
| --- | --- | --- | --- | --- | --- |
| Patient 1 | c.282+5G>A<br>(CTU2) | p.(Asp49Alafs*2)<br>(CTU2) | Fibroblast | Splice site<br>mutation<br>(SSM) | (Shaheen<br>et al., 2019) |
| Patient 2 | c.873G>A | p.(Thr247Alafs*21) | Fibroblast | Splice site<br>mutation | (Shaheen,<br>Patel, et al.,<br>2016) |
| LCLs<br>L63P<br>(0900) | c.188T>C | p.(Leu63Pro) | B-LCL | Point<br>mutation | (Shaheen<br>et al., 2019) |
| LCLs<br>SSM 1<br>(0881) | c.282+5G>A | p.(Asp49Alafs*2) | B-LCL | Splice site<br>mutation | (Shaheen<br>et al., 2019) |
| LCLs<br>SSM 2<br>(0352) | c.282+5G>A | p.(Asp49Alafs*2) | B-LCL | Splice site<br>mutation | Alkuraya<br>lab |
| LCLs<br>SSM 3<br>(0993) | c.873G>A | p.(Thr247Alafs*21) | B-LCL | Splice site<br>mutation | (Shaheen,<br>Patel, et al.,<br>2016) |
| LCLs<br>SSM 4<br>(0001) | c.873G>A | p.(Thr247Alafs*21) | B-LCL | Splice site<br>mutation | Alkuraya<br>lab, unpubl. |
| LCLs<br>DEL<br>(0260) | c.1511_1514del | p.(Leu504fs*) | B-LCL | Deletion | Alkuraya<br>lab, unpubl. |

Several patient LCLs displayed markedly reduced growth compared to controls. To test whether this defect resulted from the loss of CTU2 protein function, we transduced LCLs with retroviral particles encoding CTU2-eGFP. Wildtype CTU2 expression increased growth rates, with transduced cells outcompeting non-transduced ones in mixed cultures **(Fig. 1c, Suppl. Fig. S1c)**. This effect was pronounced in LCLs carrying splice site mutations but absent in the line with a compound heterozygous *CTU2^L63P^* mutation **(Suppl. Fig. S1c)**.

Loss of tRNA wobble thiolation causes codon-dependent ribosome pausing – particularly at CAA and AAA codons – and protein aggregation in yeast (Nedialkova & Leidel, 2015). Similar aggregation effects have been observed in human breast cancer cell lines (Rapino et al., 2021). To assess proteostasis in DREAM-PL patient cells, we analyzed protein folding, which is impacted by changes in ribosomal translation speed (Buhr et al., 2016; Nedialkova & Leidel, 2015). We used the sensor tetraphenylethene maleimide (TPE-MI), which fluoresces upon binding exposed cysteines that are normally buried in properly folded proteins and thus indicative of proteostasis defects (M. Z. Chen et al., 2017). Indeed, DREAM-PL fibroblasts exhibited significantly elevated cysteine reactivity, suggesting increased protein misfolding **(Fig. 1e).** The same was observed in two patient LCLs harboring *CTU2* splice site mutations **(Fig. 1f)**. Interestingly, all patient LCLs showed a drastic reduction in the central endoplasmic reticulum chaperone BiP (HSPA5/GRP78) **(Fig. 1g**), indicating disrupted ER homeostasis, as BiP levels would normally adjust to increased ER protein load (Bakunts et al., 2017). Consistent with altered translational control, eIF2α phosphorylation was also increased **(Fig. 1g)**, which indicates activation of the unfolded protein response. Notably, *CTU2^L63P^*LCLs showed intermediate levels of these markers, consistent with their hypomorphic growth behavior **(Suppl. Fig. S1c)**.

Collectively, DREAM-PL patient cells display altered growth and proteostasis due to impaired tRNA thiolation. Although the severity varies with cell type and the mechanism of CTU2 inactivation, the consistent reduction of thiolation across patient samples indicates a conserved negative impact at the cellular level.

### CTU2^L63P^ fails to support the formation of a functional CTU1/CTU2 complex

To obtain a mechanistic understanding of how CTU2^L63P^ leads to DREAM-PL in humans and how it compares to complete protein loss due to *CTU2* splice site mutations **(Fig. 1a)**, we assessed the functional and structural impact of this pathogenic point mutation. Leucine 63 is located at the distal end of an N-terminal α-helix, predicted by AlphaFold3 to be in contact with its substrate tRNAs **(Fig. 2a-b and Suppl. Fig. S2a, S2b)**. Substituting leucine with proline at this position is expected to act as a “helix breaker” by disrupting hydrogen bonding in the cyclic side chain (Nilsson et al., 1998). Consistent with this, the CTU2^L63P^ mutation destabilized the protein. Using insect cells for expression, wildtype CTU2 readily co-purified with CTU1, but the mutant failed to do so **(Fig. 2c).** Even when using a codon-optimized construct, CTU2^L63P^ did not form a complex with CTU1 despite being expressed, indicating that the mutation renders the protein insoluble.

**Figure 2.**
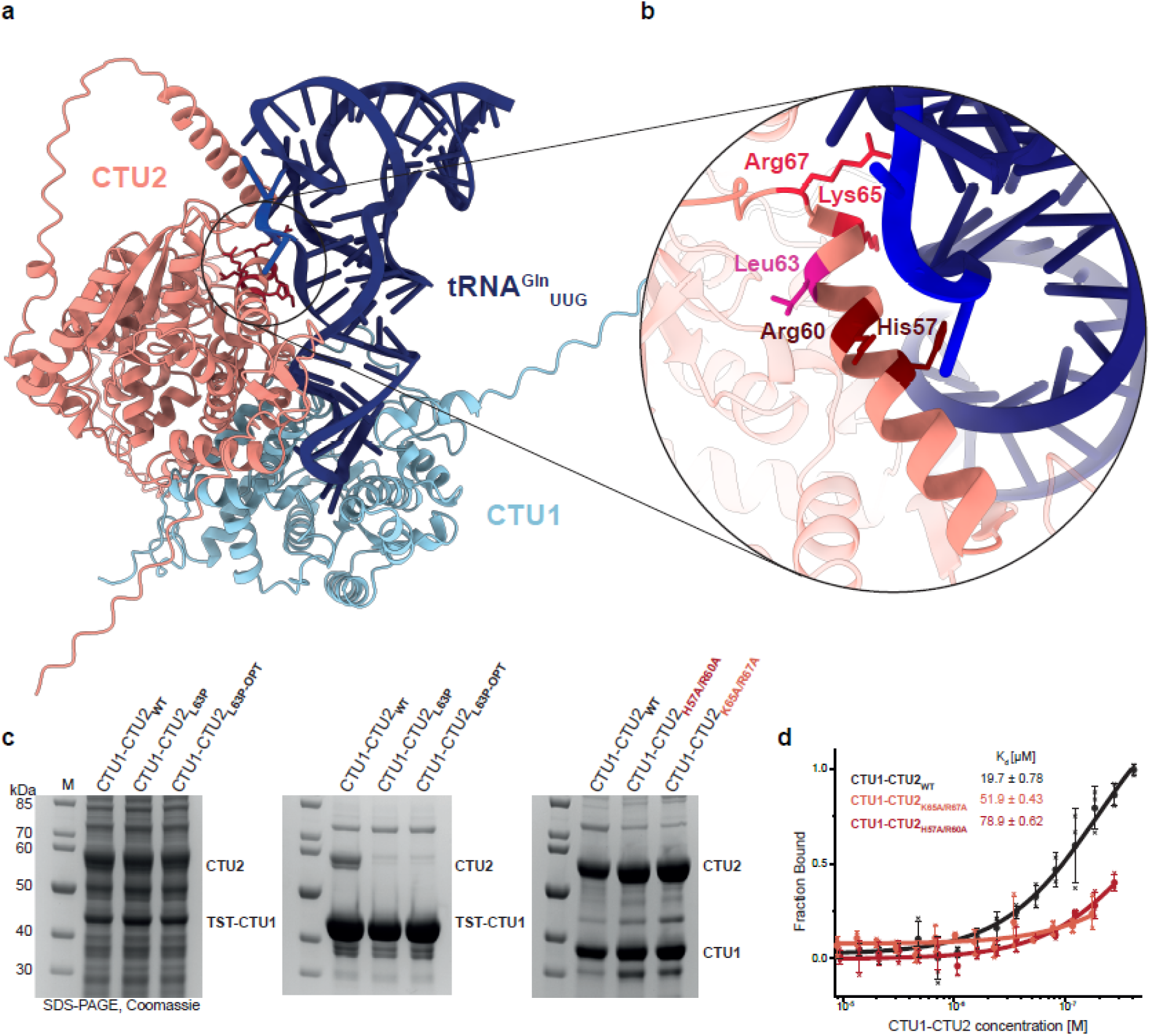
CTU2^L63P^ fails to support the formation of a functional CTU1/CTU2 complex. **(a)** Representative AlphaFold 3 model of the human CTU1:CTU2 complex bound to 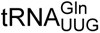. CTU1 is shown in light blue, CTU2 in red, and tRNA in navy blue. **(b)** Close-up view of the CTU2 α-helix containing Leu63, which is mutated to proline in DREAM-PL syndrome (pink). Positively charged residues selected for mutational analysis are highlighted in shades of red. **(c)** Pull-down analysis of CTU1:CTU2 complex formation and solubility. Left: whole cell extract confirming expression of CTU2^WT^, CTU2^L63P^, CTU2^L63P-OPT^ (codon-optimized for insect cell expression). Center: eluates from TST–CTU1 pull-downs assessing soluble complex formation. Right: purified human wildtype and mutant CTU1:CTU2 complexes. **(d)** tRNA-binding assay using Cy5-labeled *in-vitro* transcribed human 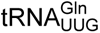 incubated with increasing concentrations of purified CTU1:CTU2 complexes. Dissociation constants K_d_ are indicated.

Interestingly, sequence analysis revealed several positively charged residues in proximity of leucine 63 – histidine 57, arginine 60, lysine 65 and arginine 76 (Fig. 2b). As these may mediate interactions with negatively charged tRNA, we generated and expressed double mutants for functional investigation. Unlike CTU2L63P, both mutants were soluble, and each stoichiometric complex could be purified in amounts comparable to the wildtype complex (Suppl. Fig. S2c). Next, we performed in vitro tRNA binding assays confirming that both CTU2H57A/R60A and CTU2K65A/R67A had markedly reduced affinity for tRNA (Fig. 2d). Together, these findings suggest that CTU2 supports tRNA binding, and that the pathogenic CTU2L63P mutation disrupts substrate, and possibly also CTU1 interactions, leading to impaired tRNA thiolation.

### Multi-omics analysis reveals transcriptome and proteome rewiring in DREAM-PL cells

To assess how cells respond to defective tRNA thiolation, we examined possible ways to adapt. Therefore, we performed transcriptome analyses in primary patient fibroblasts comparing them to healthy donor controls and found a significant upregulation of genes required for mitochondrial ATP production via oxidative phosphorylation and ribonucleotide metabolism **(Fig. 3a)**. In contrast, downregulated genes were associated with cell migration (e.g., SRC, ROCK1/2), kidney and vascular development (WNT4, SOX18), PKB/AKT survival signaling (PDGFRB, PIK3R1) and tube morphogenesis (VASP, EDNRB), in line with the developmental defects seen in DREAM-PL patients **(Fig. 3a)**.

**Figure 3.**
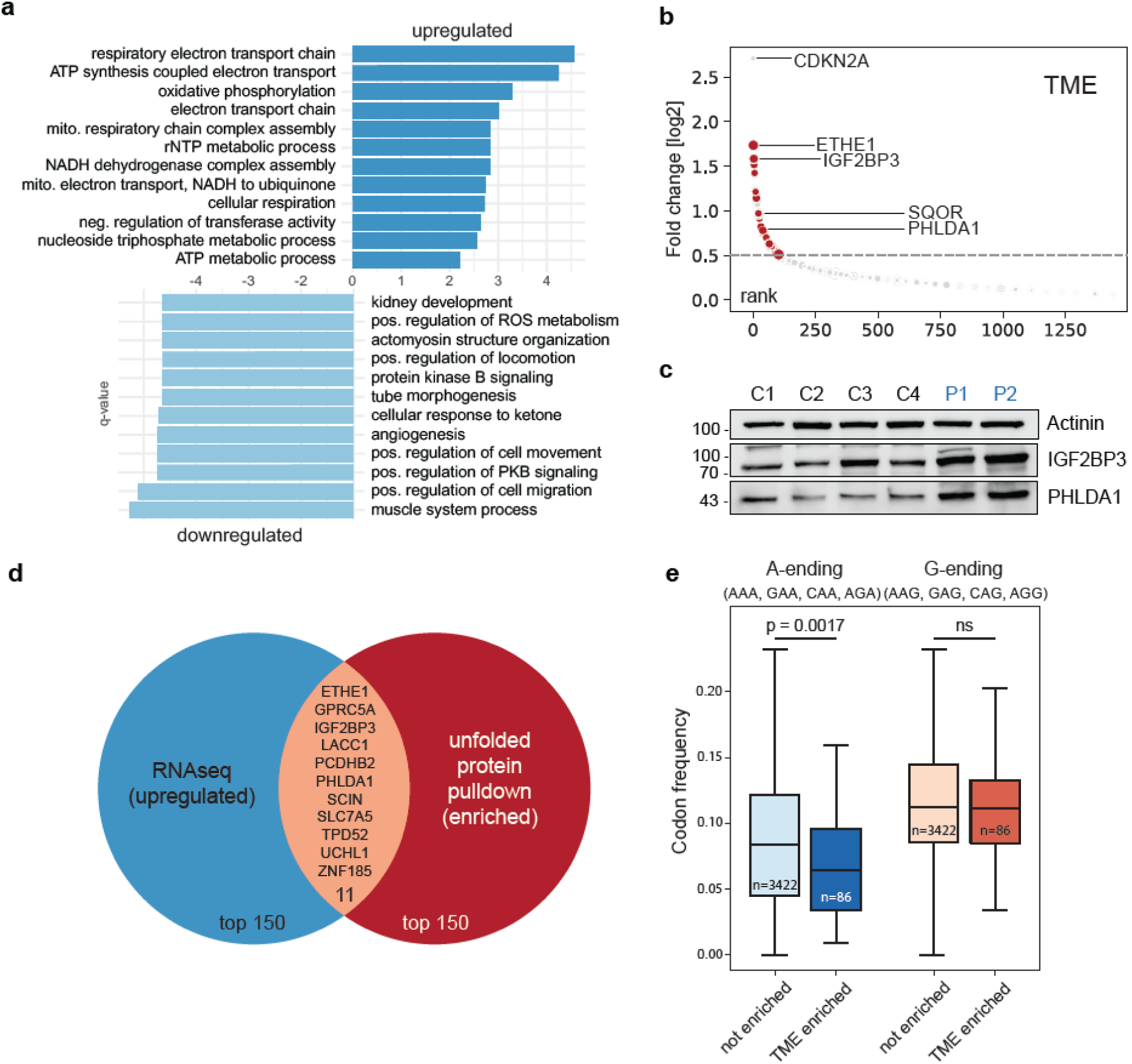
Multi-omics analysis reveals altered mitochondrial activity in DREAM-PL cells. **(a)** Gene Ontology (GO) enrichment analysis of differentially expressed genes in DREAM-PL patient fibroblasts. Bar plots show significantly upregulated (top), and downregulated (bottom) biological processes compared with healthy donor controls (n = 2 DREAM-PL fibroblast lines, n = 10 healthy donor controls; FDR < 0.05, Benjamini-Hochberg adjusted). **(b)** Unfolded proteome profiling using TME-based pulldown in patient fibroblasts. Significantly enriched proteins (adjusted p < 0.05, log_2_ fold change > 0.5) are shown in red. Dot size indicates statistical significance (–log_10_ p-value). n = 2 DREAM-PL fibroblasts profiled as technical quadruplicates; n = 4 controls profiled as technical duplicates. **(c)** Validation of selected proteins by Western blotting in patient fibroblasts. Actinin serves as loading control. **(d)** Venn diagram illustrating the overlap between upregulated transcripts and TME-enriched unfolded proteins. **(e)** Cumulative codon frequency analysis (AAA/G, CAA/G, GAA/G, AGA/G) of coding sequences of proteins identified in the unfolded proteome pulldown. n numbers are indicated. Two-tailed unpaired t-test for G-ending codons: p = 0.68.

Since CTU2-deficient cells accumulate misfolded proteins, we hypothesized that cells might transcriptionally compensate for these non-functional proteins. To test this, we profiled the unfolded proteome using TME, a second-generation TPE-MI probe **(Fig. 1e)** that selectively enriches unfolded proteins via its alkyne handle (Zhang et al., 2025). Mass spectrometry revealed an enrichment of mitochondrial proteins, oxidoreductases, and proteins involved in amino acid biosynthesis **(Fig. 3b, Suppl. Fig. S3a)**. In line with results of this STRING analysis (Szklarczyk et al., 2023), we detected significant differences in reactive oxygen species levels in patient fibroblasts compared to the healthy controls **(Suppl. Fig. S3b)**. Among the most enriched proteins were ETHE1 and SQOR, involved in mitochondrial sulfur detoxification, the m^6^A readers and mRNA-binding proteins IGF2BP1 and IGF2BP3, and factors linked to cell cycle and death regulation, including CDKN2A and PHLDA1. We confirmed expression changes of several of these candidates by immunoblotting, with IGF2BP3 and PHLDA1 showing the most pronounced increase in patient fibroblasts **(Fig. 3c)**.

Next, we performed a multi-omics analysis comparing transcriptomic and proteomic datasets and found a substantial overlap between upregulated mRNAs and TME-enriched unfolded proteins, including IGF2BP3, PHLDA1, and ETHE1 **(Fig. 3d).** This overlap indicates that signaling networks are re-wired by a cellular stress response to impaired thiolation, as many unfolded proteins are also transcriptionally upregulated in patient cells, indicative of compensatory measures that secure cell growth and survival.

Since inefficient translation of A-ending thiolation-dependent codons has been linked to protein aggregation in yeast (Nedialkova & Leidel, 2015), we assessed whether these codons are differentially represented in the coding sequences of the unfolded proteome of DREAM-PL patients. Intriguingly, proteins enriched in the TME pull-down had a lower average frequency of AAA, CAA, GAA, and AGA codons in their mRNAs compared to unchanged or underrepresented proteins, while the frequency of corresponding G-ending codons (e.g. AAG) remained unchanged **(Fig. 3e).**

### Acute CTU2 depletion induces codon-biased proteome remodeling

Because the patient cell data indicated altered proteostasis and a potential codon-specific translation defect, we generated an RNAi-based conditional *CTU2*-depletion system to directly test the consequences of timed CTU2 loss and thiolation defects in isogenic cells. Using two independent doxycycline-inducible short hairpin RNAs (shRNAs), we were able to significantly reduce CTU2 protein expression in HeLa cervical carcinoma and hTERT-immortalized retinal pigmented epithelial cells (RPE1) over time **(Fig. 4a, Suppl. Fig. S4a)**, coinciding with a significant reduction of tRNA thiolation **(Fig. 4b)**. Interestingly, CTU2 depletion markedly reduced viability in HeLa cells, while RPE1 cells showed no signs of cell death **(Fig. 4c)**. This phenotype in Hela cells was rescued by the overexpression of an shRNA-resistant variant of *CTU2*, confirming the on-target design of our construct **(Fig. 4c)**. These model cell lines recapitulated the molecular hallmarks of impaired thiolation observed in patient cells, including BiP loss and increased eIF2α phosphorylation **(Fig. 4d)**. We then asked whether proteins identified in the unfolded proteome were similarly affected upon acute *CTU2* knockdown. Indeed, ETHE1 and IGF2BP3 expression increased from day 4 onward, mirroring stress responses seen in patient cells **(Fig. 4e)**.

**Figure 4.**
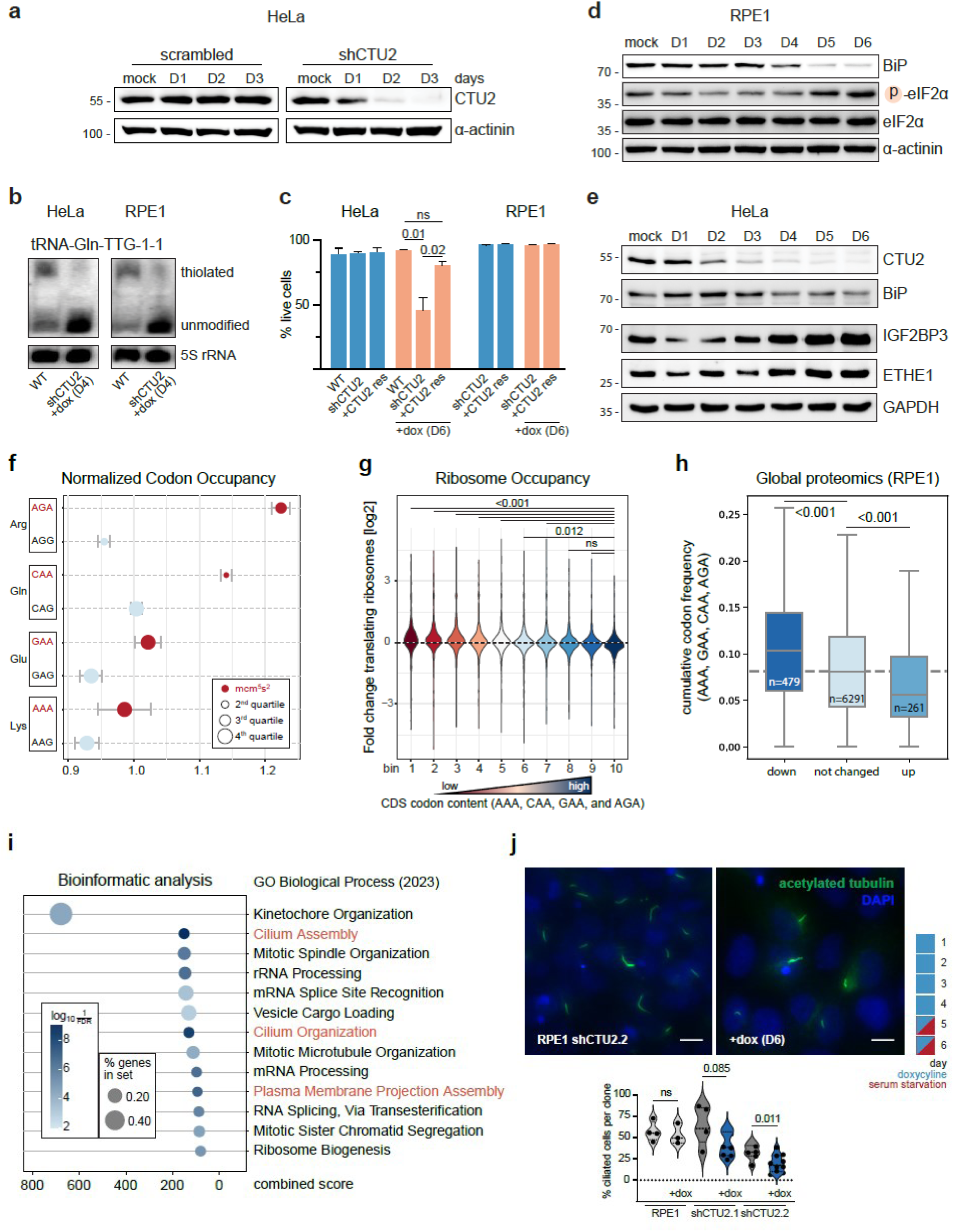
Acute CTU2 depletion induces codon-biased proteome remodeling. **(a)** Western blot analysis of doxycycline (dox)-inducible *CTU2* knockdown in HeLa cells over time. Duration of dox treatment is indicated in days. scrambled, non-targeting shRNA control. **(b)** APM-northern blot validation of tRNA thiolation loss in HeLa and RPE1 cells after 4 days of dox treatment. 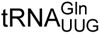 is shown representatively. **(c)** Viability analysis of *CTU2* knockdown cell lines by Annexin V staining. +CTU2 res indicates shRNA-resistant CTU2 overexpression. One-way ANOVA, WT vs. CTU2 res p = 0.264. n = 3 technical replicates. **(d)** Western blot analysis of unfolded protein response (UPR) checkpoints during progressive tRNA hypothiolation in RPE1 cells. Time points indicate days of *CTU2* depletion. **(e)** Western blot validation of selected proteins identified in DREAM-PL patient fibroblasts, analyzed under acute *CTU2* knockdown conditions. **(f)** Ribosome profiling in 96 h *CTU2*-depleted RPE1 cells compared with untreated controls. Normalized codon occupancy is shown for A- and G-ending codons of the indicated amino acids. Codons are divided into quartiles according to relative library abundance. n = 2 technical replicates. **(g)** Cumulative A-ending codon frequency (AAA, CAA, GAA, and AGA) in coding sequences (CDSs) of proteins quantified in *CTU2*-depleted RPE1 cells after 96 hours. The dashed line indicates the average A-ending codon content (7.6%) across 19,085 analyzed human CDSs. Two-tailed unpaired t-test, p > 0.001 (down and up). **(h)** Gene Ontology (GO) enrichment analysis on the top 5% of human coding sequences ranked by A-ending codon content (AAA, CAA, GAA, AGA). Enriched pathways were clustered by GO Biological Process (2023). Cilium-associated terms are highlighted in orange. **(i)** Ribosome occupancy analysis of all mRNAs measured by ribosome profiling (f). Transcripts were ranked by A-ending codon frequency (AAA, CAA, GAA and AGA) and divided into ten equal bins (bin 1 = lowest, bin 10 = highest). A negative correlation was observed between A-ending codon frequency and ribosome abundance. ANOVA with post hoc test. Adjusted p-values are indicated. **(j)** Ciliogenesis assay in *CTU2*-knockdown and wildtype RPE1 cells upon serum starvation. Representative immunofluorescence images show primary cilia (green) and nuclei (blue). The treatment scheme is shown above. Quantification of fractions of cilia-forming RPE1 cells under tRNA hypothiolation is provided. Two-tailed unpaired t-test, p-values are indicated. n = 3 technical replicates. Scale bars 10µm.

Next, to test whether thiolation loss elicits similar translational defects in human cells as reported in invertebrates (Nedialkova & Leidel, 2015; Zinshteyn & Gilbert, 2013), we performed ribosome profiling in acutely depleted RPE1 cells, a non-transformed, near-primary cell model. *CTU2* knockdown for 96 h led to a striking increase in ribosome occupancy at AGA and CAA codons **(Fig. 4f)**. Across all 61 sense codons, AGA exhibited the most pronounced increase in ribosome occupancy **(Suppl. Fig. S4b).** As expected, loss of thiolation did not affect the translation of synonymous AAG, CAG, GAG or AGG codons, which showed decreased ribosome dwell times compared to their respective A-ending codons and were comparable to fully thiolation-competent controls **(Fig. 4f)**. Given this codon-specific ribosome pausing, we next asked whether the frequency of A-ending codons of a given mRNA influences its translation under hypothiolation. Therefore, we grouped all mRNAs quantified by ribosome profiling **(Fig. 4f)** into ten bins from lowest to highest AAA, CAA, GAA, AGA content, and measured total ribosome occupancy. Strikingly, mRNAs in the highest A-ending codon bin had the fewest translating ribosomes, with a gradual and significant increase in ribosome occupancy toward bins with lower A-ending codon content **(Fig. 4g)**. These findings suggest that the loss of thiolation affects translation of A-ending codon in a dosage-dependent manner. To validate this at the protein level, we performed quantitative proteomics of *CTU2*-depleted RPE1 cells after 96 h. Consistent with the ribosome occupancy results, proteins that decreased in abundance were significantly enriched for coding sequences containing AAA, CAA, GAA, and AGA codons, whereas proteins encoded by mRNAs with few A-ending codons increased in abundance **(Fig. 4h)**.

This prompted us to perform a codon enrichment analysis to systematically identify pathways potentially disrupted in DREAM-PL syndrome. We reasoned that DREAM-PL, as a syndromic disorder, likely arises from disruption of multiple, potentially tissue-specific signaling pathways. To this end, we ranked all human coding sequences (CDS) by A-ending codon content and performed gene set enrichment analysis on the top 5% (GSEA) **(Fig. 4i)**. Strikingly, among the top 10 gene sets, three were linked to cilia biology (cilium assembly, cilium organization, and plasma membrane–bounded cell projection assembly). Additional potentially affected pathways included chromosome segregation and mitotic spindle organization, both also dependent on centrosome formation and function **(Fig. 4i)**.

Intriguingly, DREAM-PL syndrome presents with ciliopathy-like features, suggesting that the loss of tRNA thiolation may affect cilia formation or function, along with mitotic spindle performance. Given this unexpected link to ciliogenesis, we examined whether the absence of tRNA thiolation impaired primary cilia formation. Using RPE1 cells, which form primary cilia upon serum withdrawal, we compared ciliogenesis between wildtype and *CTU2*-depleted cells. Upon starvation, *CTU2* knockdown markedly reduced the fraction of cells that could form primary cilia **(Fig. 4j)**. We also observed instances of abnormal centriole counts and occasional multipolar spindles in CTU2-deficient HeLa cells, consistent with our GO-term analysis indicating defects in chromosome segregation **(Fig. 4h, Suppl. Fig. 5a-b**).

In summary, acute loss of tRNA thiolation in human cells broadly alters the proteome, with a predominant impact on pathways regulating ciliogenesis and cell division.

### *Ctu2^L63P/L63P^* knock-in mice escape DREAM-PL pathology despite reduced tRNA thiolation

CTU2 is highly conserved between mice and humans. Consistent with this, we found that the murine protein can thiolate human tRNAs, when overexpressed in *CTU2*-knockdown cell lines **(Suppl. Fig. S6a)**. Hence, we anticipated that *Ctu2^L63P^*would be equally pathogenic in mice and humans and cause comparable downstream effects.

To validate this, we used CRISPR–Cas9 to introduce the leucine-to-proline substitution into KH2 mouse embryonic stem cells for blastocyst injection, generating transgenic heterozygous animals. Surprisingly, intercrosses of *Ctu2^WT/L63P^* mice yielded homozygous *Ctu2^L63P/L63P^*offspring at only mildly reduced Mendelian ratios and a slightly skewed sex distribution **(Fig. 5a)**. Viable offspring were also born after backcrossing to the C57Bl/6N genetic background **(Suppl. Fig. S6b)**, contrasting previous reports of embryonic lethality following whole-body deletion of *Ctu1*, *Ctu2*, or *Urm1* (Birling et al., 2021; Elrick et al., 2024; Groza et al., 2023; The International Mouse Phenotyping Consortium et al., 2016). Both female and male *Ctu2^L63P/L63P^* animals were fertile and showed no developmental defects resembling those seen in DREAM-PL patients carrying the same point mutation **(Suppl. Fig. S6c)**.

**Figure 5:**
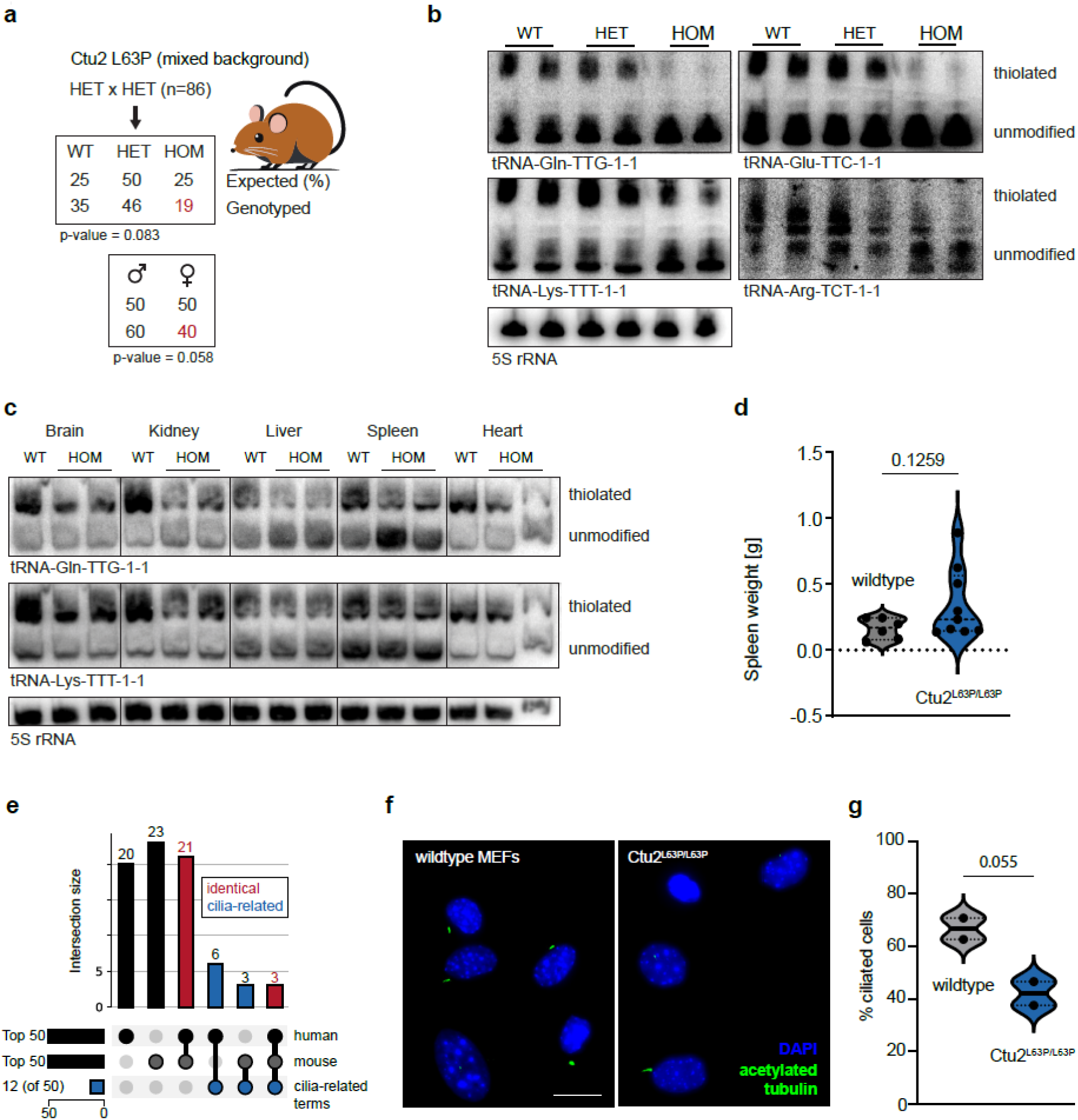
*Ctu2^L63P/L63P^* knock-in mice escape DREAM-PL pathology despite reduced tRNA thiolation. **(a)** Breeding results of heterozygous *Ctu2^WT/L63P^* knock-in mice born at Mendelian ratios, showing a trend toward underrepresentation of homozygous and female offspring. Chi-square goodness-of-fit test. p-values are indicated. **(b)** APM-northern blots detecting all thiolated tRNA species in mouse embryonic fibroblasts (MEFs). 5S rRNA serves as a loading control. **(c)** APM-northern blots for the indicated tRNA species extracted from different organs of *Ctu2^L63P/L63P^* mice. 5S rRNA serves as a loading control. **(d)** Spleen weights of aged *Ctu2^L63P/L63P^* mice (≥ 1.5 years, n = 9) compared with age-matched wildtype controls (n = 6). Two-tailed unpaired t-test. p-value is indicated. **(e)** UpSet plot showing conserved tRNA thiolation-dependent pathways between humans and mice. 24 Gene Ontology (GO) terms are conserved (red). Cilium-associated terms account for 9 (human) and 6 (mouse) GO terms, of which 3 are shared (red, right bar). Unique GO terms are shown in black.**(f, g)** Ciliogenesis assay in wildtype and *Ctu2^L63P/L63P^* MEFs. (f) Representative immunofluorescence images of primary cilia (green) and nuclei (blue). (g) Quantification of fraction of ciliated MEFs under serum starvation. Two-tailed unpaired t-test. p-value is indicated.

To confirm tRNA thiolation deficiency, we conducted APM-northern blot analyses and found a striking reduction of tRNA thiolation in mouse embryonic fibroblasts (MEFs) isolated from homozygous *Ctu2^L63P/L63P^* E14.5 embryos, highlighting the functional importance of leucine 63 both in humans and mice **(Fig. 5b)**. Furthermore, CTU2 protein levels were markedly reduced **(Suppl. Fig. S6d)**, consistent with observations in patient cells **(Fig. 1a)** and heterologous expression studies in insect cells **(Fig. 2c)**. Wildtype and heterozygous MEFs showed comparable thiolation levels, indicating that a single functional *Ctu2* allele is sufficient to maintain the thiolated tRNA pool, at least in fibroblasts. We monitored multiple tissues for tRNA thiolation and found that mutant animals showed markedly reduced thiolation across all major organs analyzed **(Fig. 5c)**. However, the ratio of thiolated to unmodified tRNAs seemed to differ among organs, with a pronounced shift best notable in the spleen **(Fig. 5c)**. Intriguingly, several aged *Ctu2^L63P/L63P^*animals analyzed presented with splenomegaly when compared to wildtype controls **(Fig. 5d**), warranting more detailed analyses of ageing phenotypes in future analyses.

Finally, we explored whether the codon biases associated with loss of tRNA thiolation are conserved among vertebrates. An analysis of coding sequences from different species with annotated human homologs showed that thiolation-dependent codon usage is most conserved in mammals and decreases progressively with evolutionary distance **(Suppl. Fig. S6e)**. Hence, we decided to make use of our mouse model and performed ribosome profiling of wildtype and homozygous *Ctu2^L63P^*MEFs and were able to confirm the pausing of ribosomes at A-ending, rather than synonymous G-ending, codons. Notably, in mice, CAA emerged as the most affected thiolation-dependent codon, in contrast to AGA in *CTU2*-depleted RPE1 cells (**Suppl. Fig. S6f, Fig. 4h**). At the pathway level, nearly half of the top AAA-, CAA-, GAA-, and AGA-enriched GO terms are shared between mice and humans, including those linked to cilia and chromosome segregation **(Fig. 5e)**. Accordingly, we repeated the ciliogenesis assay upon serum withdrawal in MEFs and found a comparable reduction of ciliation-competent cells in a state of hypo-thiolation **(Fig. 6f-g, Fig. 4j)**.

In summary, despite conserved molecular signatures of impaired tRNA thiolation, mice remain largely unaffected at the organismal level, suggesting the presence of compensatory mechanisms that allow them to escape the development of DREAM-PL–like pathology.

## DISCUSSION

Here, we investigated the cellular and organismal consequences of impaired 2-thiolation of tRNA wobble uridines in mammals, using patient-derived cells, acute cellular depletion models, and a *CTU2^L63P^* knock-in mouse. This work expands the study of tRNA thiolation from yeast to mammalian systems and provides insight into its role in the pathogenesis of DREAM-PL syndrome.

We identify CTU2 as an integral, non-catalytic component of the thiouridylase complex, likely positioning tRNA for modification by CTU1 (Fig. 2d). Interestingly, a recently characterized *S. cerevisiae* strain isolated from a patient carried an *NCS2/CTU2* gain-of-function variant (H71L) that stabilized the CTU1/2 complex to sustain tRNA thiolation and remain virulent even at elevated temperatures (Alings et al., 2023). This residue lies close to the pathogenic human L63P mutation, which we show destabilizes the CTU1/2 complex (Fig. 2c). Together, these findings illustrate that CTU2 mutations can modulate the strength and functionality of the thiouridylase complex in both directions. Notably, some bacteria and archaea perform thiolation using a single homodimeric enzyme (Shigi, 2014). Why eukaryotes evolved to use a two-subunit system remains an open question.

Once tRNA thiolation is impaired, all human cell models we profiled showed signs of impaired proteostasis, including increased protein unfolding and stress signaling, yet only some were affected in their viability (Fig. 1d-f, Fig. 4c-e). Patient and RPE1 cells tolerated hypothiolation, while HeLa cells progressively died. Transcriptomic and proteomic rewiring likely underlies this variability, enabling some cell types to adapt despite persistent proteostasis stress. A systematic follow-up on these changes (Fig. 3), especially hydrogen sulfide (H₂S) metabolism is warranted (ETHE1, SQOR), as the loss of tRNA as the final sulfur acceptor could trap sulfur at thiocarboxylated URM1, diverting it to alternative biological use (Pabis et al., 2020; Ravichandran et al., 2022; Sokołowski et al., 2024). Sensitivity to hypothiolation may also reflect changes in metabolic or translational demands, as U_34_-modifying enzymes are known to become critical under nutrient-limiting conditions or during cancer metastasis (Hermann et al., 2025; Rapino et al., 2018). Differences in RNA modification patterns could also contribute, as loss of mcm^5^s^2^ was shown to alter mRNA turnover through crosstalk with m^6^A (Linder et al., 2025).

On the translation level, impaired tRNA thiolation perturbs decoding fidelity in a codon-dependent manner. Our ribosome profiling data show markedly slowed decoding of thiolation-dependent CAA and AGA codons in mcm^5^s^2^-deficient human and murine cells (Fig. 4f, Fig. S6f), which is consistent with previously described U_34_ modification mutants (Linder et al., 2025; Nedialkova & Leidel, 2015; Zinshteyn & Gilbert, 2013). This in turn alters translational output, suggested by the underrepresentation of proteins encoded by “biased” mRNAs in the unfolded protein pulldown (Fig. 3e) and the frequency-dependent decrease of overall ribosome occupancy on these transcripts (Fig. 4i). Mechanistically, such difficult-to-translate mRNAs, together with their nascent protein products, may be degraded by cellular ribosome-associated quality control mechanisms, as shown in yeast (Wu et al., n.d.). However, codon content alone does not fully define translational efficiency. Only a subset of proteins is likely affected, as shown in yeast thiolation mutants (Rezgui et al., 2013), and additional determinants such as di-codon context, amino acid patterns and mRNA stability further modulate protein output (Linder et al., 2025; Rapino et al., 2021; Wu et al., n.d.).

Therefore, to capture a spectrum of “biased” transcripts, we performed gene ontology analysis, which revealed an enrichment for pathways related to ciliogenesis and centrosome organization (Fig. 4h). Dysfunction of cilia, cilia-anchoring centrosomes, and their coordination with the cell cycle align well with the ciliopathy-like presentation of DREAM-PL syndrome. Notably, ciliopathies are the most common cause of multisystem and renal anomalies in abnormal fetuses studied in a highly consanguineous parent cohort (Al-Hamed et al., 2022). Consistent with these observations, we confirmed reduced ciliogenesis capacity in CTU2-deficient human and mouse cells (Fig. 4j, Suppl. Fig. 5f, g).

The finding that *Ctu2^L63P/L63P^* knock-in mice exhibited strongly reduced tRNA thiolation yet completed development was unexpected (Fig. 5a–b). Together with organ-specific differences in thiolation levels (Fig. 5c), this suggest that either a low but functional threshold of tRNA thiolation is sufficient for sustaining cellular homeostasis in mice, or that individual organs differ in their dependence on thiolation for proper function. The latter is consistent with the striking tissue selectivity of DREAM-PL. Nevertheless, we observed codon-specific ribosome pausing (Fig. S6f) and reduced ciliogenesis capacity in MEFs, which may reflect a stress-dependent vulnerability, as the assay is performed under serum starvation (Fig .5f, g). Future studies employing conditional *Ctu2* ablation – like *Elp3* deletion in neural progenitors that causes microcephaly (Laguesse et al., 2015) – will be critical to define the cell type–specific requirements for tRNA thiolation and how its loss translates into organ-level pathology.

Taken together, our data support a model in which disrupted tRNA thiolation compromises translation in a context- and codon-dependent manner, specifically affecting vulnerable pathways such as ciliogenesis. Our newly established tools and models will be valuable for further mechanistic studies of DREAM-PL syndrome and may ultimately guide the development of therapies targeting epitranscriptomic dysfunction.

## Acknowledgements

We are grateful to Patrick Connolly, Tamas Szabo, and Michael J. Kraakman for insightful discussions. We thank Kaan Boztug, Georg Stary, and colleagues for providing healthy donor B-LCLs and fibroblasts, and Yuning Hong for the kind gift of TPE-MI and TME. We further thank Andrzej Chramiec-Głąbik for producing the labelled tRNA, and Julia Heppke and Mathias Drach for their contributions to histological analyses. pMSCV-IRES-GFP was a gift from William Hahn (Addgene plasmid #9044).

We thank the Molecular Discovery Platform at CeMM for assistance with proteomics analyses and the Next Generation Sequencing Platform of the University of Bern for sequencing ribosome profiling libraries.

This work was supported by the European Research Council (ERC) under the European Union’s Horizon 2020 research and innovation program (grant agreement No 101001394 to S. G.). A.V. acknowledges support by the European Research Council, ERC AdG POLICE (787171). S.A.L. was supported by Swiss National Science Foundation grants 310030_219536, and 320030_236130. This research was funded in part by the Austrian Science Fund (FWF) [10.55776/PIN1183123]. For open access purposes, the author has applied a CC BY public copyright license to any author accepted manuscript version arising from this submission.

## Author Contributions

L.E. and A.V. designed the study. L.E., M.W., C.E., F.E., V.C.S, F.M.G., Y.T., Z.Z., and N.S. performed experiments. D.M., C.E. analyzed the data. L.A. and F.S.A. provided patient samples. F.E. and T.K. established the mouse model. J.M., S.A.L., L.E, S.G. and A.V. secured funding. W.R. provided critical reagents. S.A.L., S.G. and A.V. supervised parts of the study. L.E. and A.V. wrote the manuscript with input from all coauthors.

---

**Supplementary Fig. 1.**
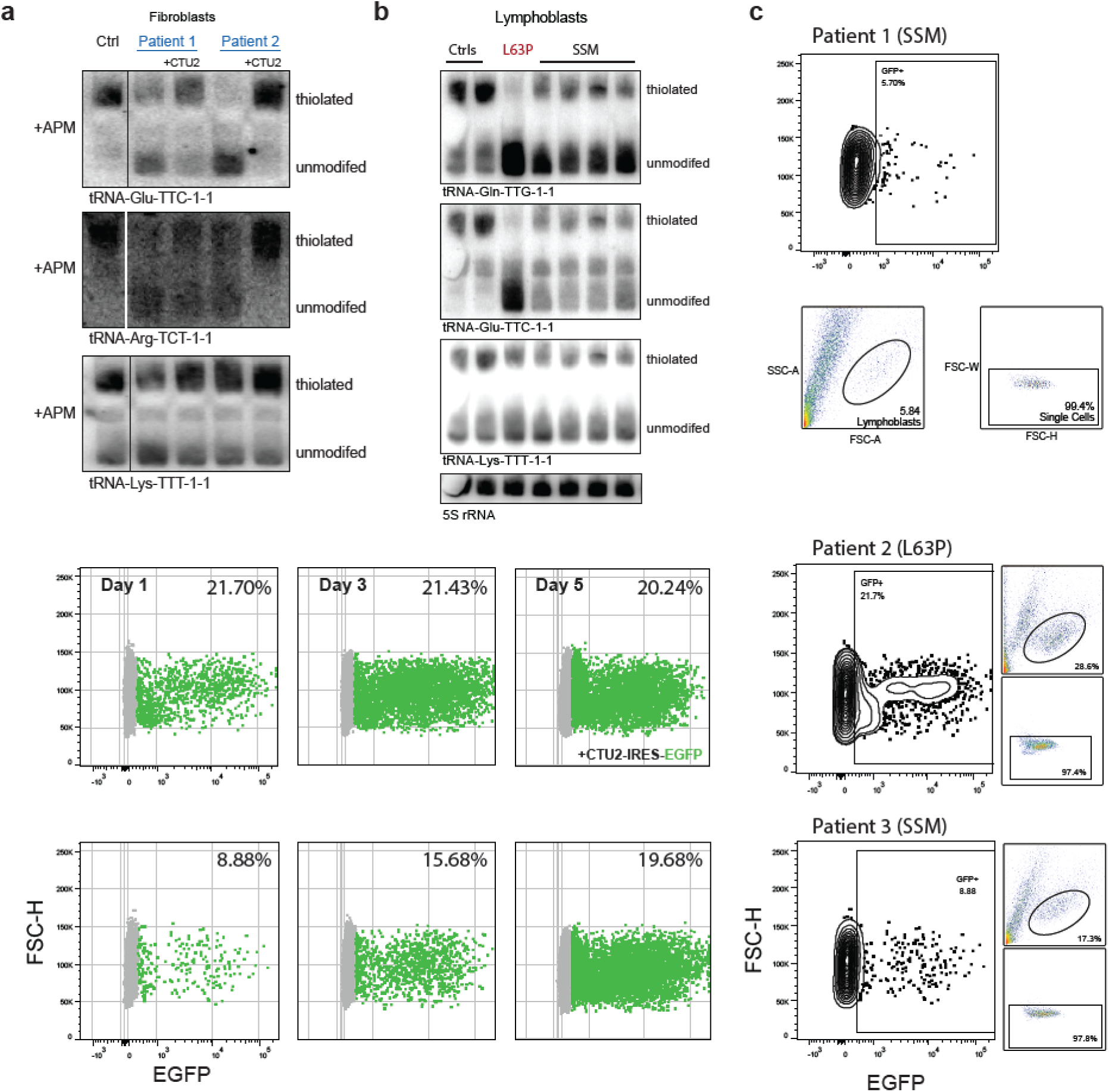
**(a)** APM-northern blot of patient fibroblasts and healthy control. The membrane was stripped and re-probed from Fig. 1b. 5S rRNA serves as a loading control. **(b)** APM-northern blot of patient lymphoblastoid cell lines (LCLs). 5S rRNA is used as loading control. **(c)** Top: flow-cytometric gating strategy for competition assays. Left: rescue of growth defects in additional patient LCLs shown in Fig. 1c. No growth advantage is observed upon CTU2 overexpression in L63P mutant LCLs. Data for patient 1 are shown in Fig. 1c. SSM, splice-site mutation; L63P, leucine-to-proline substitution at residue 63.

**Supplementary Fig. 2.**
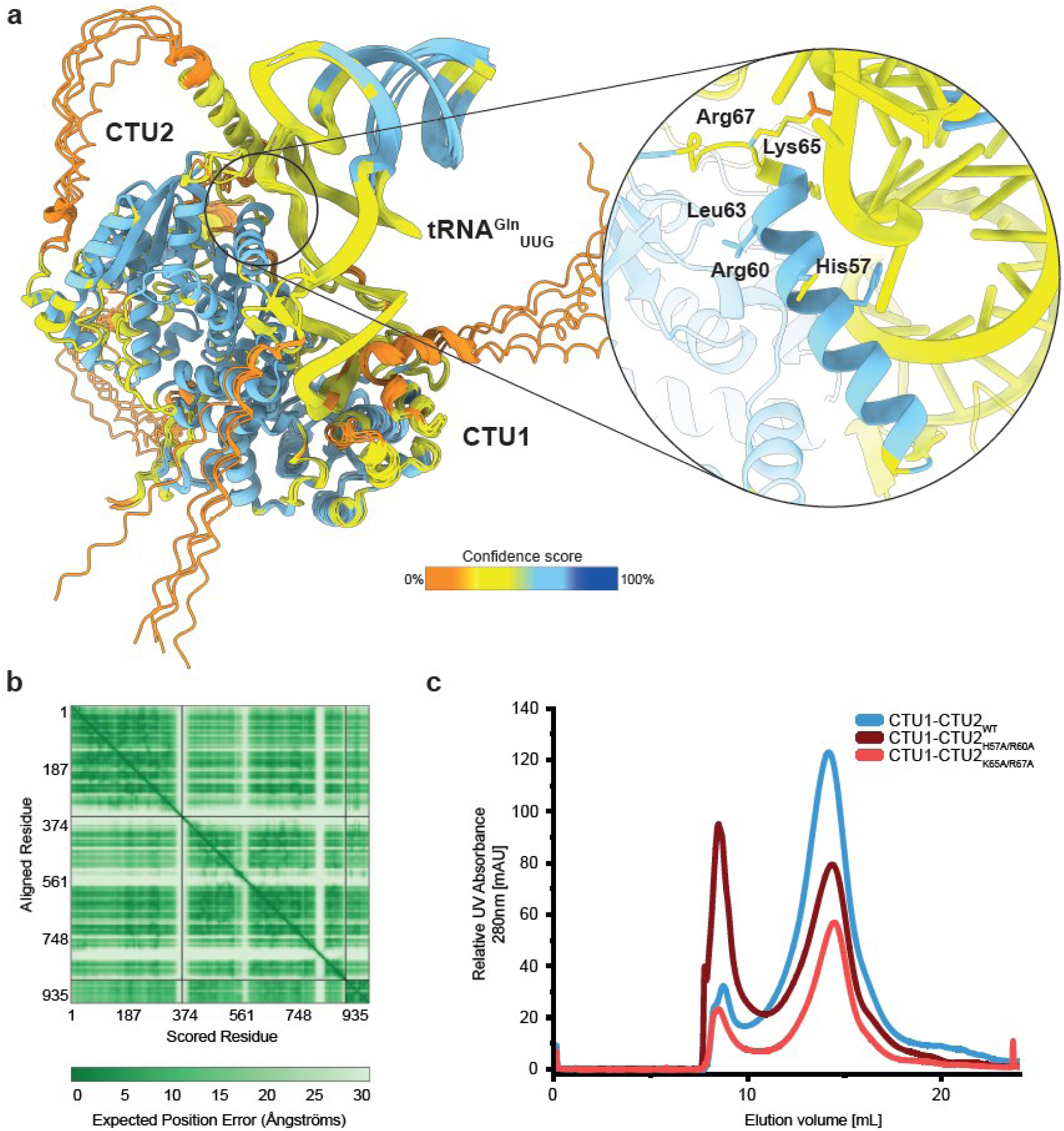
**(a)** Superimposed AlphaFold 3 predictions of the human CTU1–CTU2^WT^ complex bound to 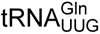, colored by per-residue confidence. Right: close-up view of the α-helix containing tRNA-binding mutants and leucine 63. **(b)** Predicted Aligned Error (PAE) plot highlighting regions of higher (dark green) and lower (pale green) inter-domain prediction confidence. **(c)** Size-exclusion chromatography profiles of purified human wildtype and mutant CTU1/CTU2 complexes from insect cells.

**Supplementary Fig. 3.**
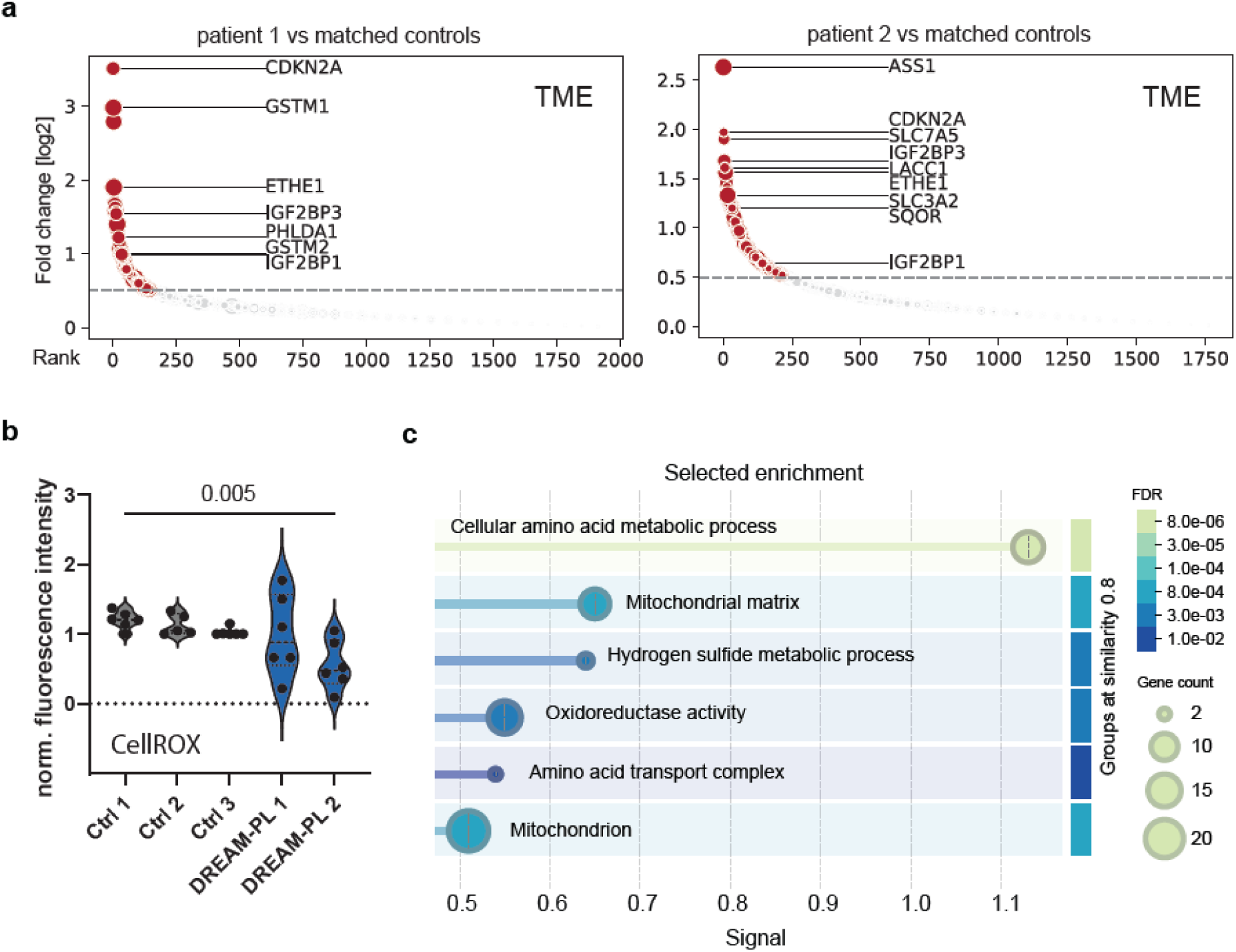
**(a)** Unfolded proteome pulldown data stratified by individual patients compared with two matched healthy controls selected by proteome similarity. Proteins detected at an adjusted p-value < 0.05 are marked in red. **(b)** Detection of reactive oxygen species in patient fibroblasts using CellROX. Welch’s t-test (Control 1 vs DREAM-PL 2) p = 0.0049. n = 3 technical replicates (measured in duplicate). **(c)** 86 proteins showing a log_2_ fold change > 0.5 in the TME-based unfolded proteome pulldown were analyzed using STRING (v12.0) without additional filtering. The top enriched Gene Ontology (GO) terms from the categories Biological Process, Molecular Function, and Cellular Component are shown. Circle size reflects the number of genes per GO term, and color indicates the false discovery rate (FDR).

**Supplementary Fig. 4.**
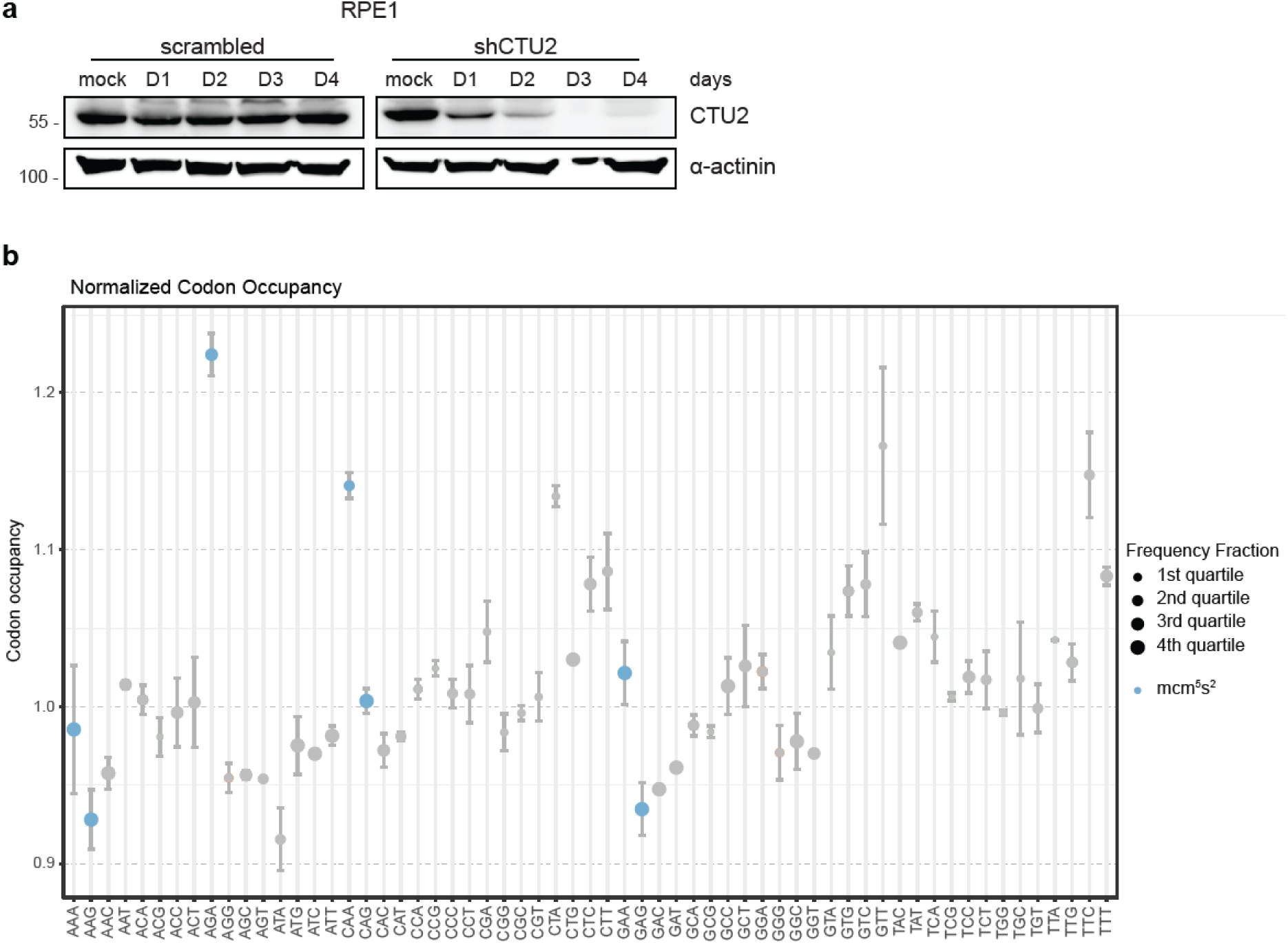
**(a)** Western blot analysis of doxycycline (dox)-inducible CTU2 knockdown in RPE1 cells over time. Duration of dox treatment is indicated in days. **(b)** Ribosome profiling in 96 h *CTU2*-depleted RPE1 cells compared with untreated controls. Normalized codon occupancy for all sense codons relative to untreated cells is shown. n = 2 technical replicates.

**Supplementary Fig. 5.**
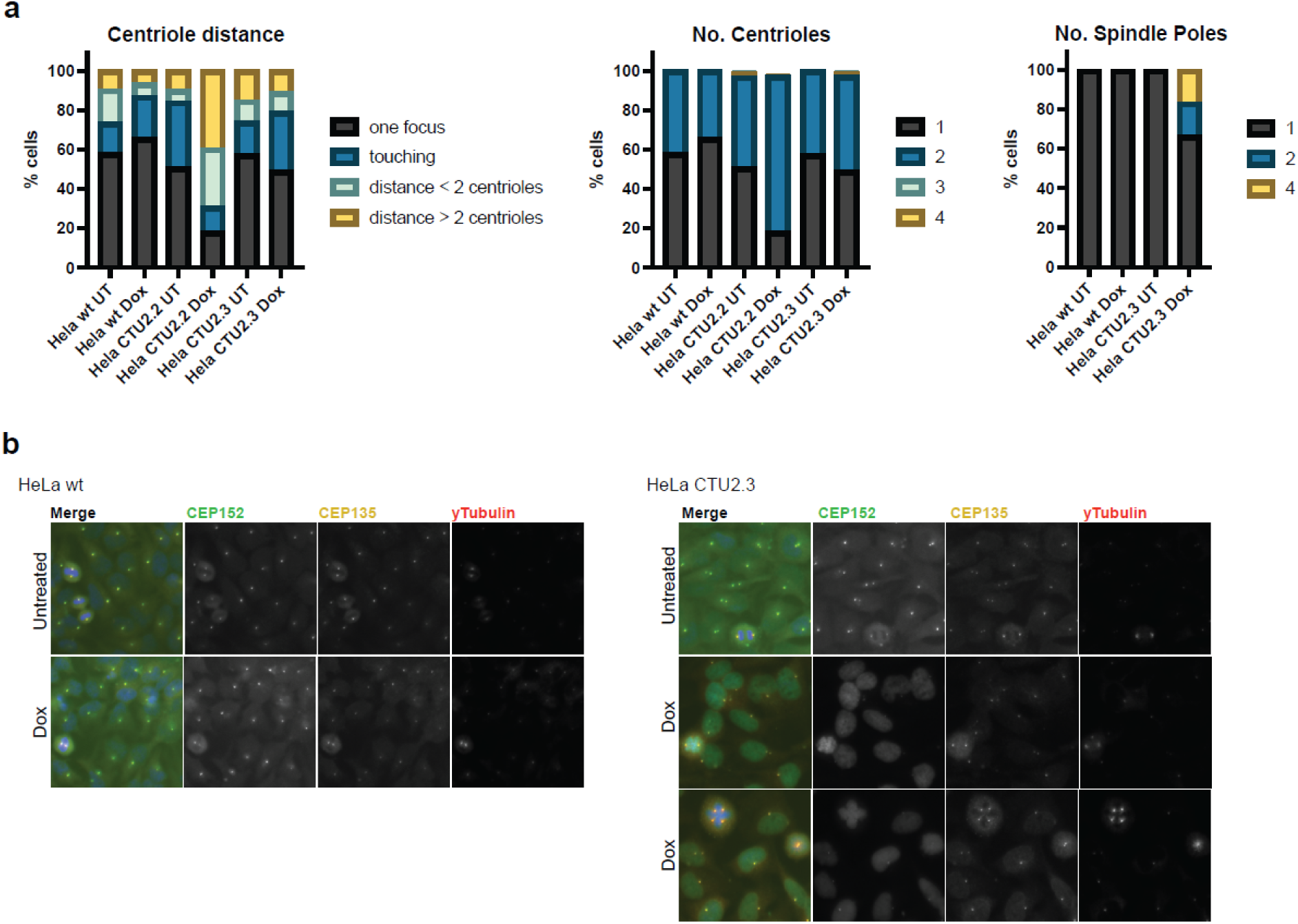
**(a)** Analysis of centriole stainings by immunofluorescence in HeLa cells upon *CTU2*-depletion. The number and distance between centrioles, as well as the number of spindle poles during mitosis, were quantified. CEP152 (green) and CEP135 (grey) mark centrioles, and γ-tubulin (red) marks spindle poles. **(b)** Representative images showing occasional multipolar spindle formation in a *CTU2*-depleted HeLa clone.

**Supplementary Fig. 6.**
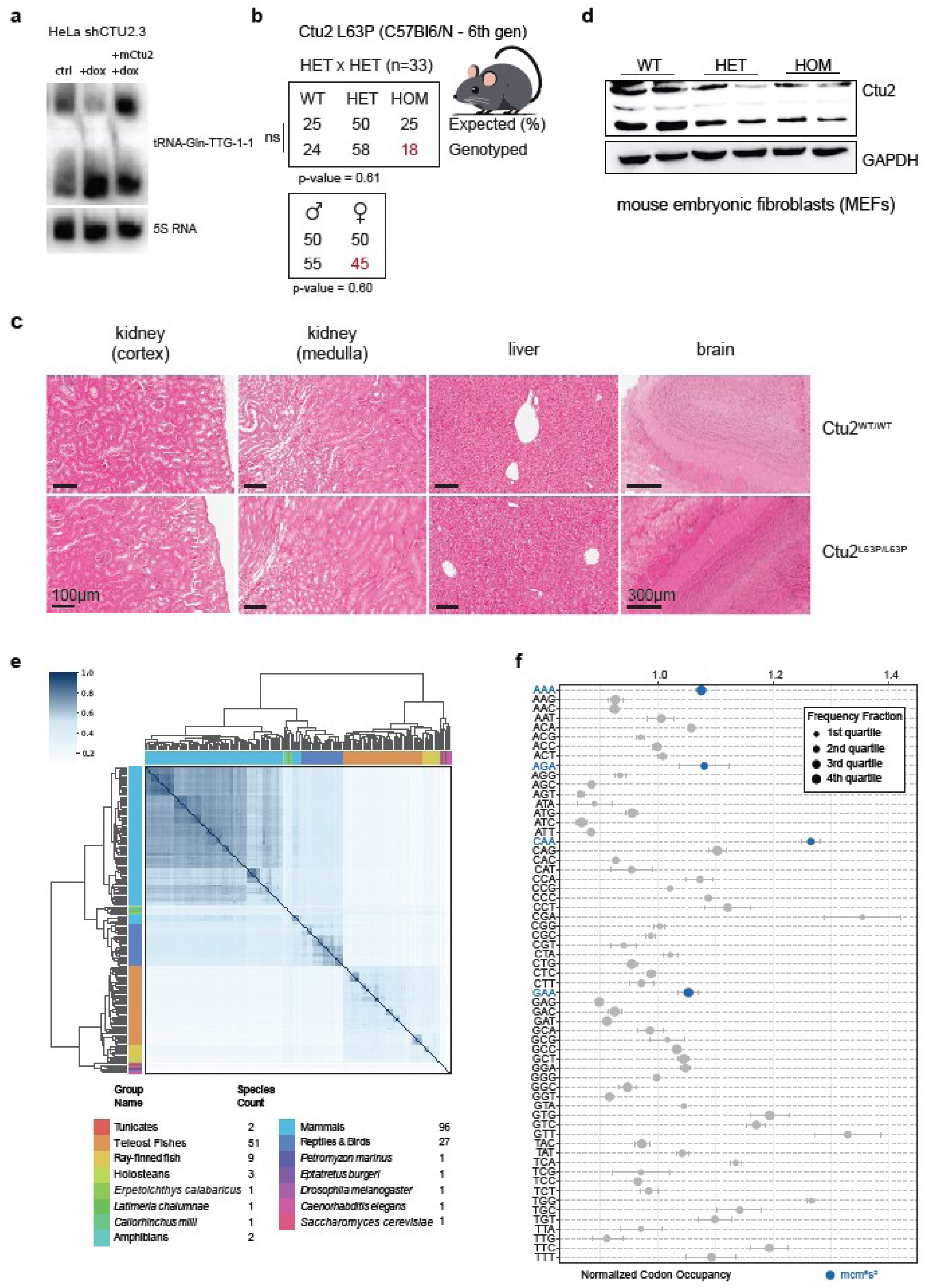
**(a)** Cellular thiolation assay using murine CTU2 protein to modify human tRNA. 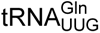 is shown representatively. 5S rRNA serves as a loading control. **(b)** Breeding results of heterozygous *Ctu2^WT/L63P^* knock-in mice backcrossed for six generations to the C57Bl/6N background, born at Mendelian ratios. Chi-square goodness-of-fit test. p-values are indicated. **(c)** Hematoxylin and eosin (H&E) staining of representative mouse organs commonly affected in DREAM-PL syndrome. Wildtype and *Ctu2^L63P/L63P^* mice are compared. Scale bars are indicated. **(d)** Representative western blot analysis of CTU2 in *Ctu2^L63P/L63P^* mouse embryonic fibroblasts (MEFs). **(e)** Cross-species comparison of A-ending codon frequency (AAA, GAA, CAA, AGA) in 197 species with annotated human homologs. Similarity scores were calculated using rank-biased overlap. Cluster group names and species counts are indicated below. **(f)** Ribosome profiling in *Ctu2^L63P/L63P^* MEFs compared with wildtype cells. Normalized codon occupancy for all sense codons relative to wildtype is shown. n = 2 biological replicates. Respective codons of mcm^5^s^2^ modified tRNAs are shown in blue.

## METHODS

### Cell culture

Fibroblasts (primary, MEFs) and cell lines (HeLa, hTERT-RPE1) were cultured in DMEM (Sigma, D5796), supplemented with 10% fetal bovine serum (Sigma, F7524) and 1% penicillin-streptomycin (Sigma, P4333). Cells were cultured at 37 °C and 5% CO_2_. STR profiling was performed for both HeLa and RPE1 (Microsynth). Cell lines were regularly tested for mycoplasma contamination by PCR.

### Western blotting

Cell pellets were lysed in RIPA buffer (50 mM Tris HCl pH8, 150 mM NaCl, 1% NP-40, 0.5% sodium deoxycholate, 0.1% SDS, 1X protease inhibitors (cOmplete EDTA-free Protease Inhibitor Cocktail, Merck). Samples were cleared and quantified using a BCA assay. 30-40µg of protein were loaded onto SDS-polyacrylamide gels (8-12%). The Mini-PROTEAN Tetra system (Bio-Rad) with Tris/Glycine/SDS buffer composition was used. A wet transfer was performed onto nitrocellulose membranes (Cytiva, GE10600002). Membranes were blocked with 5% non-fat dry milk in TBS-T. Antibodies were diluted in 1% milk in TBS-T and incubated overnight at °4C while rotating. After washing, membranes were incubated with HRP-coupled secondary antibodies (Cell Signaling, mouse 7076, rabbit 7074) for 1 hour at room temperature. ECL Detection Reagent (Cytiva, RPN2235) was used as substrate. Chemiluminescent signal was detected on a ChemiDoc MP (Bio-Rad). When required, membranes were re-probed following antibody stripping for 2× 10 mins in mild stripping buffer (0.2M Glycine, 0.1% SDS, 1% Tween20, pH 2.2), washing and blocking.

### APM-northern blotting

RNA was isolated using TRIzol. 6µg of total RNA were loaded onto 7M urea-containing 8% polyacrylamide gels supplemented with 20µg/mL APM ([(N-Acryloylamino)phenyl]mercuric Chloride (LGC Standards). RNA was transferred to positively charged nylon membranes (Hybond-N, Amersham) using a semi-dry apparatus. RNA was cross-linked to the membrane by UV light. Pre-hybridization was carried out in 6X SSC buffer with 10X Denhard’s solution, 0.5% DMSO and 100µg/mL fish sperm (Merck) for 1 hour at 40°C while rotating. Membranes were hybridized overnight with radioactively labelled oligonucleotides (ATP, [γ-32P]). Following washes with SSC buffers decreasing in concentration, the membrane was wrapped in plastic and exposed to a storage phosphor screen overnight. Signal was detected with an Amersham Typhoon laser scanner (Cytiva).

Oligonucleotide sequences for human and mouse tRNAs were taken from (Lentini et al., 2018).

### Cell line generation

For CTU2 knockdown lines we utilized the GLTR system, an all-in-one lentiviral, doxycycline inducible short hairpin RNA (shRNA) expression vector (Addgene 55790, 58246). Cloning was done as described here (Sigl et al., 2014). For the constitutive overexpression of CTU2 the retroviral vector pMIG (pMSCV-IRES-GFP) was used (Addgene 9044). Lenti/Retroviral particles production was done by co-transfection with psPAX2 (Addgene 12260 – lentiviral) / gag-pol (Addgene 35614 – retroviral) and pMD2.G (both). Spin-infection was performed at 800 x g for 90 mins at 37°C. Infected cells were selected using Puromycin (GLTR) or cells were sorted by GFP signal using a Sony SH800 sorter.

### TPE-MI staining

Cells were seeded to reach 70% confluency overnight. After trypsinization and washing with PBS, cell pellets were resuspended in 50µM TPE-MI (a kind gift from Y. Hong) diluted in 50µL PBS and incubated for 30 mins at 37 °C in the incubator. Cells were washed with PBS and analyzed with an LSR Fortessa Cell Analyzer (BD Biosciences). FCS files were analyzed using FlowJo v10.10. Statistical analysis was performed in Graphpad Prism version 10.0.3.

### AlphaFold predictions

AlphaFold 3 predictions were performed via the AlphaFold Server, using the amino acid sequences of human CTU1 (UniProt ID: Q7Z7A3), CTU2 (UniProt ID: Q2VPK5) and the sequence of tRNA^Gln^_UUG_ (GtRNAdb 2.0: chr6.trna179) at equal stoichiometric ratios. Structure predictions were carried out using default settings provided by the AlphaFold Server. The resulting models were inspected and used for visualization of the CTU1-CTU2-tRNA complex. Structural figures were prepared using ChimeraX software.

### Recombinant protein expression and purification

The codon-optimized ORFs of human CTU1 (UniProt ID: Q7Z7A3) and CTU2 (UniProt ID: Q2VPK5) were cloned into the pFASTBac vector to express proteins carrying N-terminal tags and a TEV-cleavage site to remove the tags. CTU1 was fused to 6xHis and Twin-Strep tags, whereas CTU2 was fused to a 6xHis tag, respectively. Mutated variants of CTU2 were generated by site-directed PCR mutagenesis using primers containing the desired mutation. Heterologous expression of the CTU1-CTU2 complex was performed using the Bac-to-Bac Baculovirus Expression System. In detail, the ORF-containing constructs were transformed into E. coli DH10Bac cells to generate recombinant bacmid DNA, which was subsequently transfected into Sf9 insect cells to produce recombinant baculoviruses. Individual baculoviruses and corresponding Sf9 baculovirus-infected insect cells (BiiC) were prepared separately for CTU1 and CTU2. For large-scale protein production, Hi5 cells were co-infected with CTU1 and CTU2 BiiC-derived viruses at a 1:1 virus ratio and cultured in HyClone SFM4 Insect cell medium supplemented with 0.5% fetal bovine serum (FBS) for 3 days at 27°C on a shaking platform and harvested afterwards. Cell pellets were lysed in lysis buffer (50 mM HEPES pH 7.5, 300 mM NaCl, 10% glycerol, 5 mM MgCl2, 0.1% NP-40, 1 mM TCEP supplemented with protease inhibitors cocktails and DNase I) by three freeze-thaw cycles in liquid nitrogen, followed by homogenization using an Emulsiflex C3 device (Avestin). The soluble fraction was obtained by centrifugation at 80,000 × g for 60 min. CTU1-CTU2 complexes were purified on a StrepTrap XT 5 mL affinity column (Cytiva) and eluted with elution buffer (50 mM HEPES pH 7.5, 300 mM NaCl, 5% glycerol, 1 mM TCEP, 50 mM biotin). The eluate was dialyzed overnight at 4°C against dialysis buffer (50 mM HEPES pH 7.5, 300 mM NaCl, 5% glycerol, 1 mM TCEP), using Slide-A-Lyzer dialysis cassette (Thermo Fisher Scientific). Tags were optionally cleaved with His-tagged TEV protease and removed by a second affinity purification step. Contamination and co-purifying nucleic acids were removed by using a HiTrap Heparin HP (Cytiva) affinity chromatography step. After binding the complexes were eluted with a linear gradient of heparin elution buffer (50 mM HEPES pH 7.5, 1 M KCl, 5% glycerol, 1 mM TCEP). The final eluate was concentrated using Amicon Ultra-15 centrifugal filters (30 kDa cut-off) and further separated on Superose 6 Increase 10/300 GL (Cytiva) size exclusion chromatography (SEC) column equilibrated in storage buffer (25 mM HEPES pH 7.5, 200 mM NaCl, 1 mM TCEP) to obtain pure and homogeneous protein samples. Samples were analyzed using denaturing SDS-PAGE (visualized by Coomassie Brilliant Blue) throughout all steps of the purification and complex containing SEC-fractions were pooled, concentrated, and either used for analyses or flash-frozen and stored at −80°C until further use.

### CTU2/CTU1 pull-down assays

Small scale pull-down assays were performed to evaluate expression levels, solubility, and complex formation of the human CTU1-CTU2 and mutated variants thereof. CTU1 carried an N-terminal Twin-strep tag that served as the bait protein for affinity purification of the entire complex. Hi5 insect cells (20 mL suspension cultures) were co-infected with CTU1 and CTU2 P2 baculoviruses at a 1:1 ratio and cultured under the same conditions as for large-scale expression. After three days of incubation at 27°C, cell pellets were collected and lysed by three cycles of freezing and thawing in liquid nitrogen in lysis buffer (50 mM HEPES pH 7.5, 300 mM NaCl, 10% glycerol, 5 mM MgCl_2_, 0.1% NP-40, 1 mM TCEP supplemented with protease inhibitors cocktail and DNase I). Lysates were clarified by centrifugation at 20,000 × g for 15 min, and the supernatants were incubated with 20 μL of pre-washed and equilibrated in lysis buffer Strep-Tactin Sepharose beads (IBA Lifesciences), for 2h at 4°C with gentle mixing. The beads were collected by centrifugation at 500 × g for 3 min, washed three times with lysis buffer to remove non-specific proteins and resuspended in Laemmli sample buffer. Bound proteins were released by boiling at 95°C for 5 min. Protein expression, solubility, and stoichiometry of the complexes were analyzed by SDS-PAGE on *Bolt^TM^*4-12% Bis-Tris Plus Gels (Thermo Fisher Scientific), visualized by Coomassie Brilliant Blue and documented using Chemidoc (Biorad).

### Microscale thermophoresis (MST)

Cy5-labeled, *in vitro* transcribed tRNA^Gln^_UUG_ at 50 nM was mixed with serially diluted purified human CTU1-CTU2 complex and mutants thereof in MST buffer (25 mM HEPES pH 7.5, 150 mM NaCl, 1 mM TCEP, 2 mM MgCl_2_, 0.025% Tween-20) and incubated at 20°C for 30 min. Samples were loaded into premium capillaries (MO-K025, NanoTemper Technologies) and measured using the Monolith Pico instrument (NanoTemper Technologies) equipped with MO.Control software (NanoTemper Technologies). Data from at least three independent experiments (n≥3) were analyzed with MO.Affinity Analysis software (NanoTemper Technologies), and K_d_ values were determined based on the fitted, averaged binding curves. Graphs were plotted in OriginPro software (OriginLab).

### Transcriptomics

5×10^5^ human dermal fibroblasts, from DREAM-PL patients or sex-matched healthy donor controls, were seeded into 10 cm dishes. After 48 hours, RNA was extracted from cultured fibroblasts using a kit according to manufacturer recommendations (Monarch Total RNA Miniprep Kit, New England Biolabs). RNA was polyA enriched and samples prepared for multiplexed. The library was sequenced on a NovaSeq X (Illumina) using paired end sequencing on an S4 flow cell with 300 cycles. Following demultiplexing, differential gene expression analysis using DEseq2 (1.18.1) was performed.

Gene Ontology (GO) enrichment analysis for biological processes was conducted with clusterProfiler (v3.6.0) using the human annotation database org.Hs.eg.db (v3.5.0). Significantly enriched GO terms were identified based on an FDR < 0.05, and visualization was performed using clusterProfiler’s built-in plotting functions.

### Unfolded proteome identification

5×10^5^ fibroblasts were seeded into 10cm dishes. After 24 hours, cells were harvested and transferred in PBS to 2 mL microcentrifuge tubes. Cells were collected, resuspended in 50µM TME in 300µL cysteine-free DMEM (Gibco, 21013024) and incubated for 30 mins at 37°C and 5% CO2. Cells were washed twice with PBS. Pellets were snap frozen in liquid nitrogen and stored at −80°C. Cell pellets were lysed for 30 mins at 37°C in lysis buffer (1x PBS, 1% SDS - Sigma, 71736, 2mM MgCl2 - Invitrogen, AM9530G, 1x Protease inhibitors - Thermo, 78437, 1x Benzonase - Merck, 170746) and sonicated at 4°C (Bioruptor Pico, Diagenode, 90 sec ON – 30 sec OFF, 4 cycles). Lysates were cleared by centrifugation for 15 mins at 18.000 x g. Supernatants were transferred to lo-bind tubes (Eppendorf) and quantified (Thermo, 22660).

800µg of protein in 150µL lysis buffer were used for the Click reaction and further diluted with 450µL of potassium phosphate buffer (61.34mM K2HPO4*3H2O - Sigma, P5504, 38.21mM KH2PO4 - Sigma, P0662). Click reagents were added sequentially – biotin-PEG3-azide, freshly 1:2 mixed CuSO4 - Sigma, C1297/ THPTA - Sigma, 762342, aminoguanidine HCl - Sigma, 396494, and sodium ascorbate - Roth, 3419.1 (final concentrations: 170µM, 230µM/1.15mM, 5mM, 5mM). The reaction was incubated for 1 hour at 25°C while rotating. Proteins were precipitated with ice cold acetone. The air-dried protein pellets were resuspended in 1% SDS and resolubilized using an ultrasonic bath (Diagenode). Proteins were reduced by 4.5mM TCEP at 56°C for 1 hour while rotating, adjusted to a neutral pH and alkylated at 25°C for 30 mins using iodoacetamide. Enrichment of biotinylated proteins was done with PBS-equilibrated streptavidin-agarose beads (Thermo, 20353). Enrichment was carried out at 25°C for 1 hour while rotating. The beads were washed 16x using 8M urea in PBS. Beads were washed first with PBS then resuspended and washed with digestion buffer (50mM ammonium bicarbonate, 0.2M guanidine hydrochloride, 1mM calcium chloride, HPLC grade ddH2O). 1µg trypsin was added to each sample and left for 16 hours for off-bead digestion.

Peptides were harvested and purified using solid phase extraction. CPA and channel normalization was performed. Samples were labelled using isobaric TMTpro 16plex labels (Thermo Fisher) and pooled. On-tip high pH (10) fractionation was performed (5 fractions). 10µL were injected per faction on an UltiMate 3000 RSLCnano system coupled to Orbitrap Fusion Lumos (Thermo Fisher), 5 x 190 min run, OTITOT (SPS-MS3 quantification). Data analysis was performed using Proteome Discoverer 2.4 SP1.

### Detection of ROS levels

The detection of ROS levels in fibroblasts was done according to manufacturer instructions (CellROX Deep Red Flow Cytometry Assay Kit, Invitrogen, C10491). Briefly, 1×10^5^ fibroblasts were seeded to reach 70% confluency. Cells were harvested with trypsin, transferred to 1.5 mL microcentrifuge tubes, washed with PBS and resuspended in 250 µL of complete DMEM. CellROX Deep Red reagent was diluted 1:10 in DMSO and 1 µL added to each sample at a final concentration of 1µM and incubated for 30 mins at 37°C. After 15 mins 1 µL of 1:4 diluted Blue Dead Cell stain solution in PBS was added for the remaining 15 mins. Samples were acquired on an LSR Fortessa Cell Analyzer (BD Biosciences). FCS files were analyzed using FlowJo v10.10. Statistical analysis was performed in Graphpad Prism version 10.0.3.

### Annexin V

Cells were harvested and washed with PBS at relevant timepoints. Cell pellets were resuspended in 100 µL Annexin V Binding Buffer (422201, BioLegend). Samples were stained with 5 µL of FITC Annexin V antibody (640906, BioLegend) and 10 µL of propidium iodide for live cell gating (0.5 mg/mL) for 15 mins at RT in the dark. 400 µL of Annexin V Binding Buffer was added and samples were acquired on an LSR Fortessa Cell Analyzer (BD Biosciences). FCS files were analyzed using FlowJo v10.10. Statistical analysis was performed in Graphpad Prism version 10.0.3.

### Ribosome profiling

Libraries were prepared according to (Kim et. al., 2021) with minor modifications. Briefly, 3 OD_260_ U of crude cell lysate were digested in 700µl of lysis buffer with 150U RNAse I (AM2294, Invitrogen) for 1 h at 22 °C and 1400 rpm. The reaction was stopped with 5 µl of SUPERaseIn (AM2696, Invitrogen). To purify the digested monosomes, samples were separated by ultracentrifuging the extracts in a 10-50% sucrose gradient in a SW41 rotor at 35000 g for 3 h at 4 °C, and further fractionated using a Piston Gradient Fractionator (Biocomp). Monosome fractions were collected and RNA extracted from the sucrose using hot phenol/chloroform and separated by a 15 % TBE-urea gel. Subsequently ribosome protected fragments were excised from the gel. The fragments were dephosphorylated and ligated to an adaptor carrying 6 randomized positions prior to being reverse transcribed using a primer with 3 randomized positions. After cDNA synthesis, rRNA was depleted using a custom oligo pool. Finally, the depleted cDNA was circularized, and libraries were PCR-amplified and PAGE purified. Library quantification and sequencing were performed by the next generation sequencing platform of the University of Bern.

Sequencing data from ribosome profiling was pre-processed by removing the adapter and randomized nucleotides (three at the 5’ end and 6 at the 3’ end) using the FASTX_toolkit (Hannon, 2010). Non-coding reads were removed by mapping against non-coding RNA, including rRNAs, tRNAs, snoRNAs and other annotated lncRNAs using Bowtie (v1.2.3) (Langmead et al., 2009). The retained reads were mapped to all annotated open reading frames in humans (GRCh38) with unique mapping mode allowing at most one nucleotide mismatch using the parameters “-v 1 -m 1 –norc –best –strata”. Codon translation speed was calculated as in (Nedialkova & Leidel, 2015) by normalizing the frequency of the A site codons to that in the +1, +2, +3 sites, followed by calculating the ratio of the translation rate between mutant and wild type. To focus the analysis on translation elongation, 15 codons were excluded at both end of the CDS.

### Proteomics

#### Sample preparation - FASP + C18 cleanup

Cells were lysed in 0.4 mL lysis buffer (2% SDS, 50 mM HEPES, 1 mM PMFS, 1x protease inhibitor cocktail from Roche). Samples were resuspended and incubated at room temperature for 20 minutes before they were sonicated in a Branson ultrasonic processor with micro tip on ice (0.5 seconds on / 0.5 seconds off, 30 seconds total, 20% input). The lysate was centrifuged at 16,000 g for 10 minutes at 20°C and supernatants were transferred to a fresh tube. The protein concentration was measured with Pierce BCA Protein Assay (23227) according to manufacturer’s instructions. The FASP protocol was employed for proteolytic digestion of 100 µg protein according to (Wiśniewski et al., 2009) with slight modifications. Shortly, DTT (83.3 mM) was added to the lysis buffer and incubated at 95°C for 5 minutes. Microcon 30 Ultracel YM-30 filter units were primed once with 200 µL urea buffer (8 M urea in 100 mM Tris/HCl, pH 8.5). Samples (100 µg) were loaded onto the filter units at 14,000 rcf at 20°C for 15 minutes, washed with UA solution (8 M Urea in 100 mM Tris/HCl, pH8.5). Alkylation was performed with 200 µl 50 mM Iodoacetamide in UA solution. Samples were again centrifuged (14,000 rcf, 10 minutes) after an incubation time of 30-minute incubation in the dark. Next, samples were washed twice with 100 µl UA and then twice with 100 µl 50 mM TEAB. Samples were then digested in 40 µl 50 mM TEAB, pH8.5, at a protein to enzyme ratio of 50:1. The spin columns were sealed and incubated at 37 °C overnight. On the next day, samples were spun down (14,000 rcf, 20 minutes) and the columns were washed once with 40 µL 50 mM TEAB and once with 50 µL 0.5M NaCl. The combined filtrate was acidified with 30% TFA until a pH below 3 was achieved. Samples were then subjected to C18 cleanup with PierceTM Peptide Desalting spin columns according to manufacturer’s instructions (ThermoFisher Scientific; 89851, 89852).

#### Peptide quantification

Desalted peptides were reconstituted in LC-grade water, and peptide concentrations were determined using the Pierce™ Quantitative Fluorometric Peptide Assay (Thermo Fisher Scientific, 23290) according to manufacturer’s instructions. Fluorescence measurements were performed on a SpectraMax® i3x Multi-Mode Microplate Reader (Molecular Devices).

#### TMT labeling

Peptides were subsequently buffered to a final concentration of 50 mM HEPES, pH 8.5. Labeling was performed at a peptide concentration of ≥1 µg/µL using TMT reagent prepared in anhydrous acetonitrile at 16.7 µg/µL, applied at a TMT-to-peptide mass ratio of approximately 2:1 to 3:1. The final acetonitrile concentration during the labeling reaction was 35% (v/v). Samples were incubated for 1 hour at room temperature with gentle agitation (∼600 rpm). Labeling reactions were subsequently quenched by the addition of 5% hydroxylamine solution in HEPES buffer to achieve a final hydroxylamine concentration of 0.2–0.4%, followed by incubation for 15 minutes at room temperature. Labeled peptides were pooled, dried by vacuum centrifugation, and reconstituted in 0.1% trifluoroacetic acid for subsequent processing. Labeling efficiency was routinely assessed following final analysis.

#### Offline fractionation of TMT pool

Pooled TMT labelled sample was concentrated and desalted by using a Pierce Peptide Desalting Spin Columns (89851 and 89852) according to manufacturer’s instructions with minor modifications: peptide elution was done in 2 steps: 40% ACN, 100mM TEAB and 70% ACN, 100 mM TEAB solution. Eluates were pooled and brought to dryness in a speedvac vacuum concentrator. Dry peptides were resuspended in 30ul 10mM ammonium formate (pH 10) and fractionated by reverse-phase chromatography at basic pH by using a Gemini-NX C18 (150 × 2 mm, 3 μm, 110 A,) column (Phenomenex, Torrance, USA) on an Ultimate 3000 RSLC micro system (Thermo Fisher Scientific) equipped with a fraction collector. Peptides were separated at a flow rate of 50 µl/min in 10 mM ammonia formate buffer (pH 10.0) and eluted over a 70 min nonlinear gradient from 0 to 100% solvent B (90% acetonitrile, 10 mM ammonium formate, pH 10.0). Thirty-six concatenated fractions were collected in a time-based manner (at 30 s intervals) between minute 11.5 and 57. Fractions were immediately acidified by adding 5ul 30% TFA. After fractionation organics were removed in a vacuum concentrator and sample were stored at −20C until MS analysis.

#### LC-ESI-MS/MS data acquisition

For MS analysis fractions were reconstituted in 150 ul 0.1% TFA and 5 µl were injected.

For proteomic data acquisition, a nanoflow LC−ESI-MS/MS setup comprised of a Dionex Ultimate 3000 RSLCnano system coupled to a Fusion Lumos mass spectrometer (both ThermoFisher Scientific) was used in positive ionization mode. MS data acquisition was performed in data-dependent acquisition (DDA) mode. For proteome analyses, peptides were delivered to a trap column (Acclaim™ PepMap™ 100 C18, 3 μm, 5 × 0.3 mm, Thermo Fisher Scientific) at a flowrate of 5 μL/min in HPLC grade water with 0.1% (v/v) TFA. After 10 min of loading, peptides were transferred to an analytical column (ReproSil Pur C18-AQ, 3 μm, Dr. Maisch, 500 mm × 75 μm, self-packed) and separated using a stepped gradient from minute 11 at 8% solvent B (0.4% (v/v) FA in 90% ACN) to minute 61 at 28% solvent B and minute 81 at 40% solvent B at 300 nL/min flow rate. The nano-LC solvent A was 0.4% (v/v) FA HPLC-grade water. MS1 spectra were recorded at a resolution of 60,000 using an automatic gain control target value of 4 × 10^5 and a maximum injection time of 50 ms. The cycle time was set to 2 seconds. Only precursors with charge state 2 to 6 which fall in a mass range between 360 to 1300 Da were selected and dynamic exclusion of 30 s was enabled. Peptide fragmentation was performed using higher energy collision dissociation (HCD) and a normalized collision energy of 35%. The precursor isolation window width was set to 1.3 m/z. MS2 spectra were acquired at a resolution of 30,000 with an automatic gain control target value of 5 × 10^4 and a maximum injection time of 54 ms. Using synchronous precursor selection, the top 10 fragment ions of the MS2 scans were isolated and subjected to HCD fragmentation in the linear ion trap using 55% normalized collision energy. MS3 spectra were acquired in the Orbitrap at a resolution of 50,000 over a scan range of 100 to 1000 Th, using an AGC target of 1e5 and a maximum injection time of 120 ms. The reagent tag type in the filer IsobaricTagExclusion was set to TMTpro.

#### Data analysis

For all DDA measurements, MaxQuant (version 2.6.7.0) with its built-in search engine Andromeda was used for peptide identification and quantification (Cox et al., 2011; Tyanova, Temu, & Cox, 2016). MS2 spectra were searched against all Swiss-Prot canonical protein sequences obtained from UniProt (UP000005640, downloaded: 19 March 2025), supplemented with common contaminants (built-in option in MaxQuant). Trypsin/P was specified as the proteolytic enzyme. Precursor tolerance was set to 4.5 ppm, and fragment ion tolerance to 20 ppm. The minimal peptide length was defined as seven amino acids, and the “match-between-run” function was disabled. Quantification was performed on the MS3 level. “18plex” (TMTpro) was selected as isobaric labels and the reporter mass tolerance was set to 0.003 Da. For proteome analyses, carbamidomethylated cysteine was set as fixed modification and oxidation of methionine and N-terminal protein acetylation as variable modifications. The FDR was set to 100%. These search results were then used as input files for oktoberfest (v.0.8.3) (Picciani et al., 2024). We performed Prosit rescoring and quantification via the picked-group-FDR approach (v.0.8.1) (Gessulat et al., 2019; The et al., 2022). The Prosit models Prosit_2020_irt_TMT for retention time prediction and Prosit_2020_intensity_TMT for intensity prediction were employed. The output files were filtered at 1% FDR.

Perseus was used for data analysis (Tyanova, Temu, Sinitcyn, et al., 2016). Briefly, “common contaminants”, “reversed”, and “only identified by site” were filtered out, intensities were log2-transformed, and median-centric normalized. Samples were then categorically annotated and the “Hawaii plot” function with default values was performed. The obtained matrix was exported for further investigations.

### Codon bias analysis

For the codon-bias analysis we ranked all human coding sequences (CDS) based on their content of AAA, GAA, CAA and AGA codons. In brief, we first downloaded a set of CDS nucleotide sequences from Ensembl (release 113) and subsequently counted the occurrences of all codons and computed the codon percentage for each CDS (occurrence(codon) / sum(occurrences(CDS))). We then used the resulting codon frequencies to rank all CDS according to the sum of AAA, GAA, CAA and AGA codon frequencies and computed enrichments on the top 5% of the ranked CDS with gseapy’s Enrichr API (Fang et al., 2023).

### Codon bias conservation analysis

All Ensembl release 113 species that had annotations of genes and their human homologs were compared with respect to their A-ending codons of interest AAA, GAA, CAA, AGA. In brief, we downloaded coding sequences (CDS) for 197 of the 310 species present in the Ensembl database release 113 and annotated the respective genes for their human homologs via Biomart using the API provided by gseapy 1.1.3. Only CDS with an annotated human homolog were retained. We then computed the percentage of A-ending codons of interest (AAA, GAA, CAA, AGA) for each coding sequence of each gene in each species. Subsequently, we utilized these data to compute species similarities by calculating the rank biased overlap of the top 10% highest ranking genes for each species (*i.e.* the top 10% of genes with the highest A-ending codon of interest content) implemented in rbo 0.1.3 (https://github.com/changyaochen/rbo). We then augmented this similarity matrix by clustering the accompanying Ensembl species tree with the UPGMA algorithm after pruning the tree to only contain the compared species. Finally, we plotted the resulting similarity matrix together with its species cluster annotation as a clustermap (*i.e*. rows and columns are clustered by UPGMA) using seaborn’s clustermap implementation (version 0.13.2).

### Immunofluorescence

Cells were grown on coverslips and fixed in 1.5% paraformaldehyde and then in −20°C cold methanol for 4 minutes each. The samples were blocked for 1 hour in 2.5% FBS, 200 mM glycine, and 0.1% Triton X-100 in PBS and incubated with the respective primary antibodies in the same buffer for 1 hour. After washing, the cells were incubated with secondary antibodies (Invitrogen) and DAPI, washed and mounted in Prolong Gold Antifade (Life Technologies, P36930). Primary antibodies used were CEP135 (rabbit polyclonal, Alexa 555-conjugated, 1:500, (LoMastro et al., 2022)), γ-Tubulin (goat polyclonal, Alexa 647-conjugated, 1:500, (Levine et al., 2017)), CEP152 (rabbit polyclonal, Alexa 488-conjugated, 1:500, (LoMastro et al., 2022)), Acetyl-α-Tubulin (Rabbit polyclonal, Lys40 (D20G3) Cell Signaling 5335T, 1:1000). Cells were imaged using a DMi8 inverted microscope (Leica Microsystems) or a Thunder Imager (Leica Microsystems) with an Olympus 63× 1.42 NA oil objective.

### Generation of Ctu2 L63P mice

The Ctu2^L63P^ mouse strain was generated by mutating KH2 mouse embryonic stem cells *in vitro* using CRISPR/Cas9, followed by their injection into blastocysts. In brief, KH2 cells were co-electroporated with the pSpCas9(BB)-2A-GFP (PX458) vector (Addgene 48138) (Ran et al., 2013) encoding a guide RNA targeting the required locus (sgRNA: 5’GACCCGGTTCTTCCCAAGCA3’) and a complementary ssDNA oligo repair template (ssDNA: 5’ACAGGGTTTGTTTCAAGGCATTCTACGTTCACAAATTCcGgGCtATGCcTGGcAAGAAtCGGGTCATCTTTCCTGGGGAGAAGGTACGATGTGTGGT 3’, with lowercase bases indicating mutations) (Bio-Rad GenePulse Xcell Electroporator). The repair template contained the mutation of interest and silent mutations for genotyping. GFP positive cells were sorted one day after electroporation (BD FACSAria™ III) and used to generate single cell clones. Clones were screened by PCR and validated by Sanger sequencing, before being injected into blastocysts and implanted into surrogate mothers. Germline transmission was confirmed by genotyping PCR (frw: 5’GCACGCCCCTTTTCC3’, rev: 5’gCCAgGCATaGCcCg3’) and Sanger sequencing.

